# Structural and functional vascular dysfunction within brain metastases is linked to pembrolizumab inefficacy

**DOI:** 10.1101/2023.08.25.554868

**Authors:** Albert E. Kim, Kevin W. Lou, Anita Giobbie-Hurder, Ken Chang, Mishka Gidwani, Katharina Hoebel, Jay B. Patel, Mason C. Cleveland, Praveer Singh, Christopher P. Bridge, Syed Rakin Ahmed, Benjamin A. Bearce, William Liu, Elies Fuster-Garcia, Eudocia Q. Lee, Nancy U. Lin, Beth Overmoyer, Patrick Y. Wen, Lakshmi Nayak, Justine V. Cohen, Jorg Dietrich, April Eichler, Rebecca Heist, Ian Krop, Donald Lawrence, Jennifer Ligibel, Sara Tolaney, Erica Mayer, Eric Winer, Carmen M. Perrino, Elizabeth J. Summers, Maura Mahar, Kevin Oh, Helen A. Shih, Daniel P. Cahill, Bruce R. Rosen, Yi-Fen Yen, Jayashree Kalpathy-Cramer, Maria Martinez-Lage, Ryan J. Sullivan, Priscilla K. Brastianos, Kyrre E. Emblem, Elizabeth R. Gerstner

## Abstract

Structurally and functionally aberrant vasculature is a hallmark of tumor angiogenesis and treatment resistance. Given the synergistic link between aberrant tumor vasculature and immunosuppression, we analyzed perfusion MRI for 44 patients with brain metastases (BM) undergoing treatment with pembrolizumab. To date, vascular-immune communication, or the relationship between immune checkpoint inhibitor (ICI) efficacy and vascular architecture, has not been well-characterized in human imaging studies. We found that ICI-responsive BM possessed a structurally balanced vascular makeup, which was linked to improved vascular efficiency and an immune-stimulatory microenvironment. In contrast, ICI-resistant BM were characterized by a lack of immune cell infiltration and a highly aberrant vasculature dominated by large-caliber vessels. Peri-tumor region analysis revealed early functional changes predictive of ICI resistance before radiographic evidence on conventional MRI. This study was one of the largest functional imaging studies for BM and establishes a foundation for functional studies that illuminate the mechanisms linking patterns of vascular architecture with immunosuppression, as targeting these aspects of cancer biology may serve as the basis for future combination treatments.

## Introduction

A feared complication of solid tumor malignancies is the development of brain metastases (BM) due to limited CNS penetrant treatments. Immune checkpoint inhibitors (ICI) have emerged as a promising strategy, as these agents exhibit intracranial efficacy for melanoma and lung cancer. However, investigation of ICI in BM of non-melanoma or lung histologies has been limited. We recently published a phase 2 study that demonstrated 42.1% of enrolled patients, spanning diverse tumor types, derived intracranial benefit from pembrolizumab. In addition, we unexpectedly observed durable intracranial efficacy for a subset of patients, many possessing tumor types that do not traditionally respond to ICI (e.g. breast, sarcoma)^1^. To build upon these promising treatments, there is a critical need to non-invasively interrogate ICI’s effects on the BM tumor microenvironment (TME), as it is not feasible to obtain serial biopsies to understand why some patients benefit and others experience only toxicity with no response. Identifying biological differences that underlie ICI response and resistance may shed light on potential mechanisms of intracranial efficacy to ICI.

ICI resistance has been linked to facets of a dynamic TME that promote immune evasion^2,3^. Tumor vascular physiology is a facet of the TME that is closely linked to, and may modulate, immunogenicity of the TME. Structurally and functionally aberrant vasculature, characterized by dysregulated angiogenesis and hypoxia, promotes recruitment of regulatory T cells^4–6^, reduces the cytotoxic activity of dendritic and effector T cells^7,8^, and hampers trafficking of immune cells^9,10^. Furthermore, the mechanical stress exerted on the blood vessels from macroscopic tumors promote release of pro-angiogenic factors that polarizes macrophages to an immunosuppressive M2 phenotype and inhibits dendritic cell maturation^11–13^. Vascular endothelial growth factor inhibition prunes immature vessels and reverses immunosuppression by augmenting effector T cell infiltration and function^3,14,15^, suggesting that vascular normalization is linked to improved immunosurveillance and treatment response. Therefore, studying and targeting abnormal tumor vasculature may have therapeutic potential.

To date, longitudinal structural and functional vascular changes within tumors, and its implications for immune function, have not clearly been defined in human studies. Given the unique physiology of meningeal lymphatics^2,3^ and biological differences between intracranial and extracranial metastases^16^, mechanisms of immune evasion within the central nervous system likely differ from those in the extracranial space and remain poorly understood. In clinical practice, there is frequently discordant response between BM and patient-matched extracranial tumors, where extracranial metastases will respond but a BM will progress^1,17–19^. Therefore, we sought to quantify longitudinal vascular changes within BM in the hopes of identifying mechanisms of ICI efficacy and vascular phenotypes associated with ICI response or resistance.

Dynamic susceptibility contrast (DSC) and dynamic contrast enhanced (DCE)-MRI shed light on dynamic vascular changes by non-invasively measuring relative cerebral blood volume (CBV), cerebral blood flow (CBF)^20^, and vascular permeability (K^trans^)^21^, which reflect different aspects of structural and functional vascular biology for gliomas and BM^20,22–26^. Therefore, by providing detailed insights in vascular phenotypes, a cardinal facet of tumor biology, perfusion imaging addresses inherent limitations in standard T1-weighted post-contrast imaging. For example, DSC and DCE-MRI analysis revealed vascular patterns within the tumor and peri-tumor areas that differentiate between radio-necrosis and tumor progression, which often have similar radiographic appearances on post-contrast imaging^26–30^. In addition, perfusion imaging may have promise as a predictive biomarker in ICI response. Exploratory DCE-MRI analysis in a cohort of 15 extracranial melanoma patients being treated with ICI revealed an increase in apparent diffusivity and decreased tissue heterogeneity, as measured by diffusion kurtosis, associated with ICI response^31^.

Using a cohort of 44 BM patients treated with pembrolizumab in a phase 2 trial (NCT02886585)^1^, we combined analysis of dual-echo DSC and DCE MRI with histopathology of BM tissue to understand how vascular-immune cross-talk contributes to ICI efficacy. As prior research has demonstrated a robust link between vascular architecture, vascular efficiency, and immune phenotypes^32–34^, we hypothesized that ICI is most effective in cancers possessing a structurally balanced vascular network that facilitates an immune-stimulatory TME and minimizes the immunosuppressive impact of hypoxia^2,3^. To test this theory, we used Vessel Architectural Imaging (VAI) to quantify detailed structural and functional facets of vascular physiology, such as vessel size index^22^. VAI’s performance as a quantitative metric for *in vivo* tumor vascular morphology has been validated in prior work^22,35–37^.

## Results

### Patients

Of the 79 enrolled patients with parenchymal BM, 58 patients obtained dual-echo DSC and DCE MRI. Of these patients, 44 had evaluable MRI data. Reasons for exclusion included: lost to follow-up after one cycle of pembrolizumab (N=6), measurable disease on the MPRAGE sequence but not within the FOV for the DSC sequence (N=6), or motion artifact (N=2; **Figure 1**).

**Figure 1:**
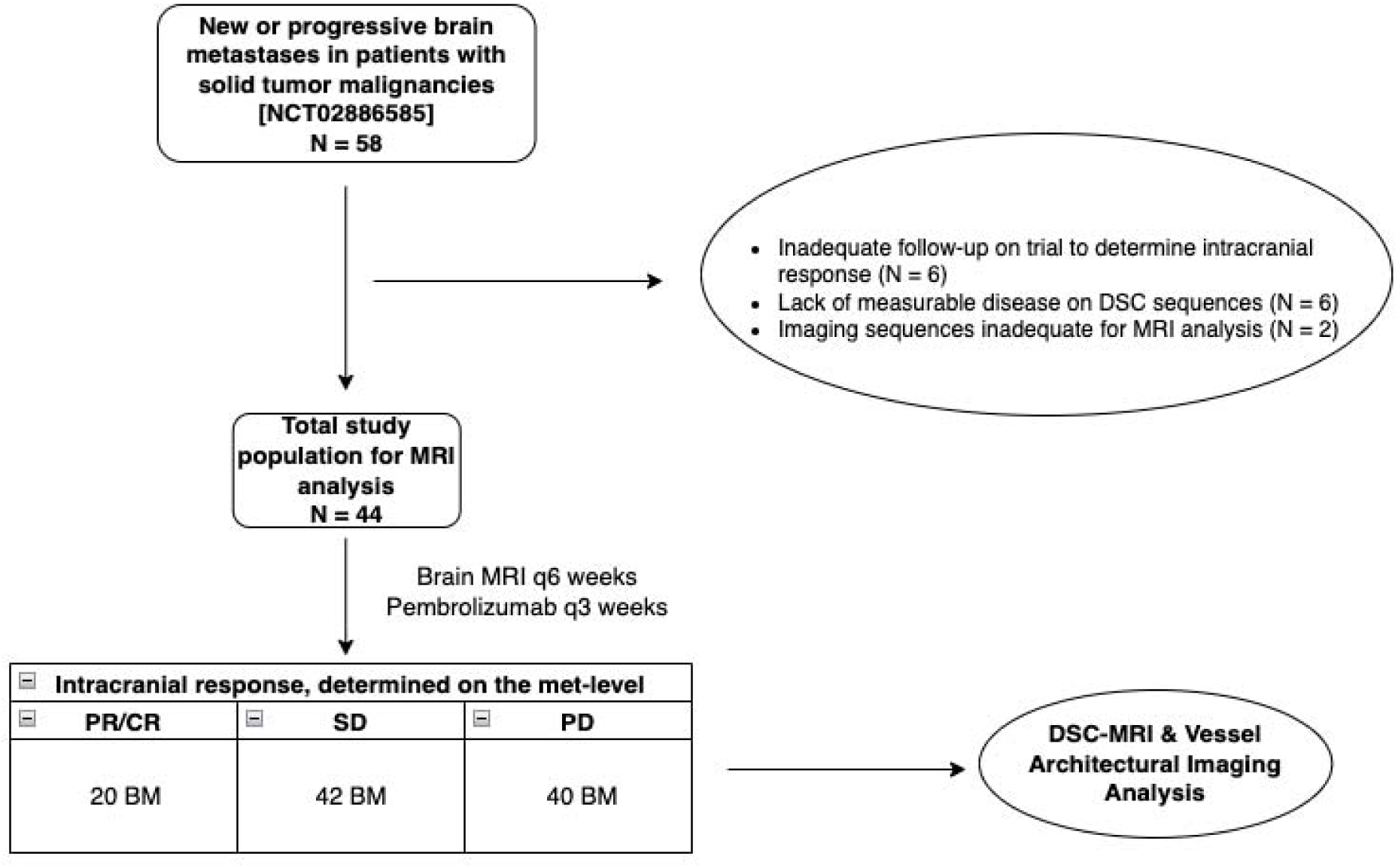
Study schema. 58 enrolled patients, while on treatment with pembrolizumab, obtained perfusion MRI due to available funding. 44 patients had evaluable MRI data. Intracranial response was defined on the metastasis-level using Response Assessment for Neuro-Oncology (RANO) criteria. These brain metastases were then used in Vessel Architectural Imaging analysis.

After MR processing, 102 individual BM within our cohort of 44 patients were suitable for analysis. Of these 102 BM, the most common histologies included breast (N=58), melanoma (N=7), NSCLC (N=7), and pituitary (N=5). Histology of other BM, demographic information, and prior treatment status are listed in **Table 1**. Per RANO-BM, 2 BM were categorized as complete response (CR), 18 as partial response (PR), 42 as stable disease (SD), and 40 as progressive disease (PD). Given the small number of CR BM, PR and CR BM were grouped together for analysis. As only 2 of the 20 PR/CR BM met the SNR voxel cutoff for analysis at or after the second post-treatment visit, we restricted our analysis to a comparison of the pre-treatment and the 6-8 weeks post-treatment MRI scans. The vascular status of all analyzed BM before and after treatment with pembrolizumab is summarized in **Supplemental Table 1**. Examples of acquired parametric images, post contrast MRI images, hysteresis plots, and vortex directions for three BM are provided in **Supplemental Figures 1-3**.

**Table 1:**
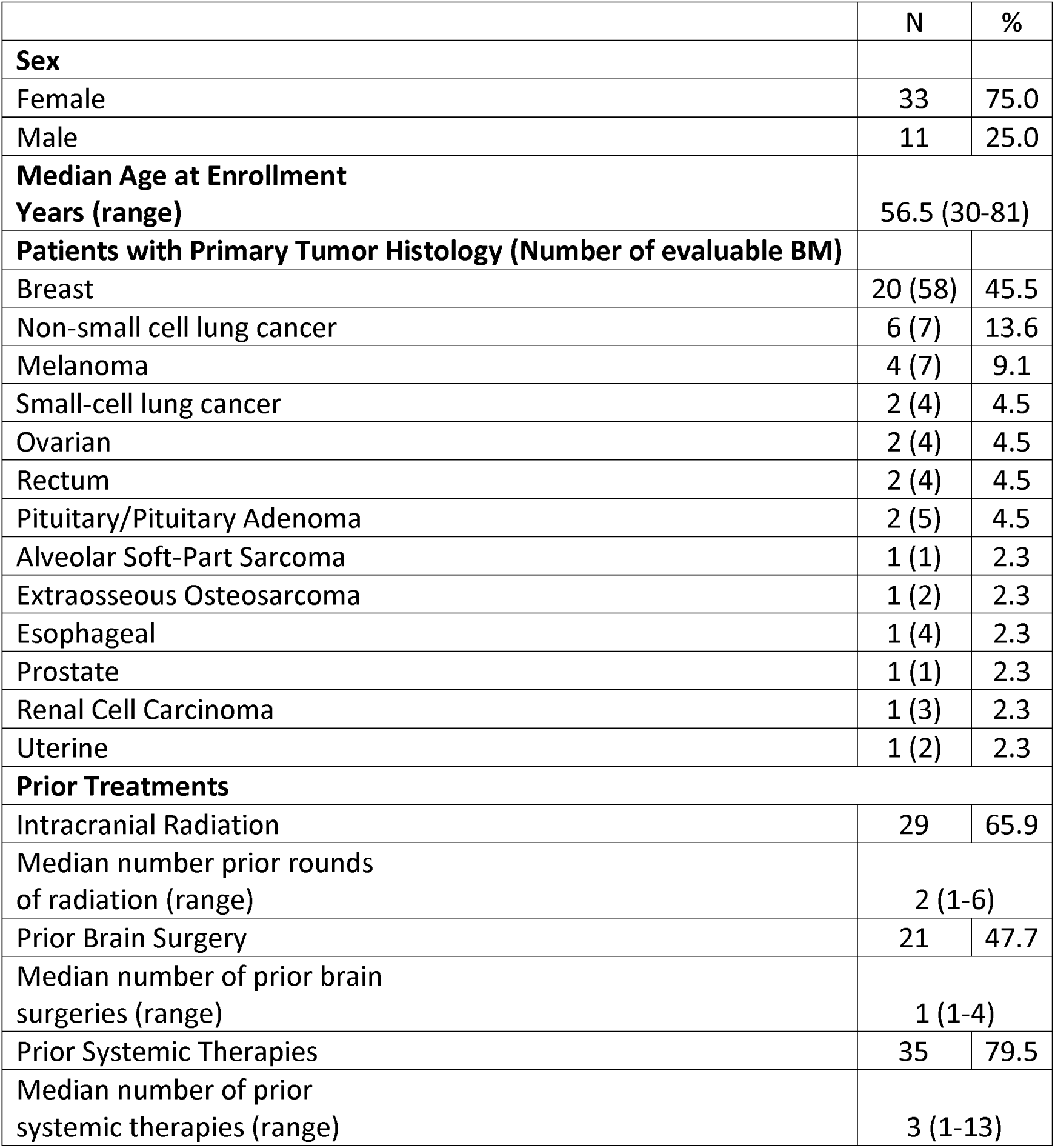
Patient demographics and disease characteristics at enrollment.

### A structurally balanced vascular network within BM is linked to an immunogenic microenvironment and favorable ICI response

When assessing the vascular network, we found that BM responding to ICI possessed a higher degree of small-caliber vessels than did BM with SD or PD (**Figure 2** – left panel) before and after treatment with pembrolizumab. We observed a low fraction of microvasculature associated with PD BM, suggesting that these BM possess a structurally unbalanced vascular TME dominated by larger-caliber vessels. Consistent with these findings, median vessel sizes were larger post-treatment for BM with SD (p=0.0004) and PD (p=0.002), compared to PR/CR (**Figure 3**). Additionally, vessel size increased longitudinally during treatment with pembrolizumab for BM with SD (p=0.0006) and PD (p=0.02), whereas no significant change was noted for PR. To corroborate the observation that the unbalanced vascular architecture of ICI-resistant BM is due to distension of both small and large vessels, no longitudinal differences were noted in GE or SE CBV with respect to intracranial response (**Supplemental Figure 4**). A lack of change in pan-vascular (GE) and microvascular (SE) blood volumes indicate that a portion of vessels of all calibers are enlarged without additional vessel recruitment. Taken together, these findings suggest that a certain structural balance of small and large-caliber vessels within tumors is ‘therapeutically favorable’ and thus facilitates ICI efficacy (**Figure 4**).

**Figure 2:**
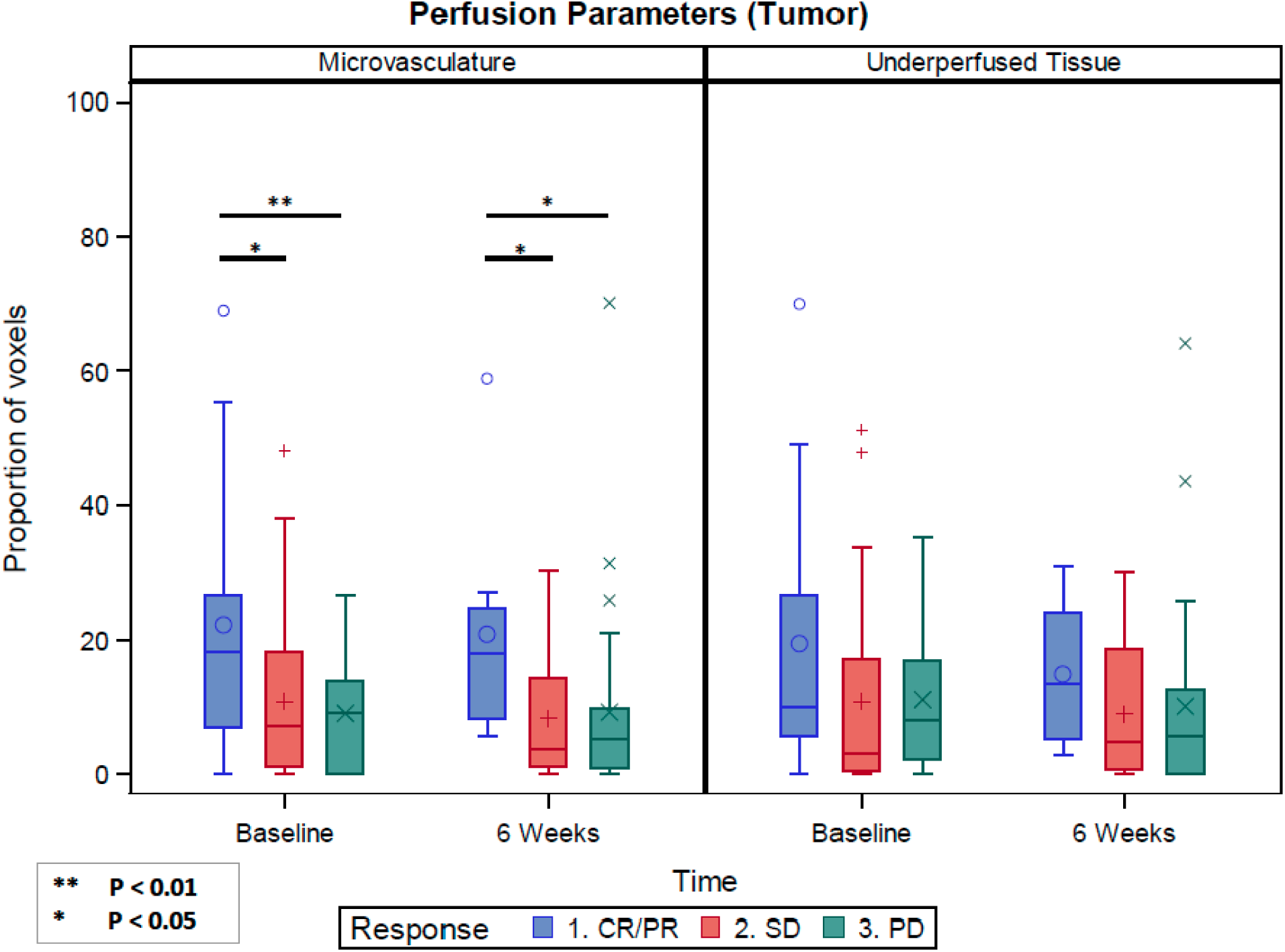
Intracranial response is associated with a structurally balanced vascular architecture. BM that responded to ICI had a significantly higher proportion of microvasculature within the contrast-enhancing area of the tumor, compared to SD or PD for both pre-treatment (p=0.04; p=0.007, respectively) and post-treatment (p=0.01; p=0.04). Similarly, PR/CR BM displayed a trend towards a longitudinal decrease in under-perfused tissue (p=0.13). In addition, BM that responded to ICI had a non-significant trend towards a higher degree of under-perfusion compared to SD or PD for both pre-treatment (p=0.06, p =0.10) and post-treatment (p=0.08, p=0.08).

**Figure 3:**
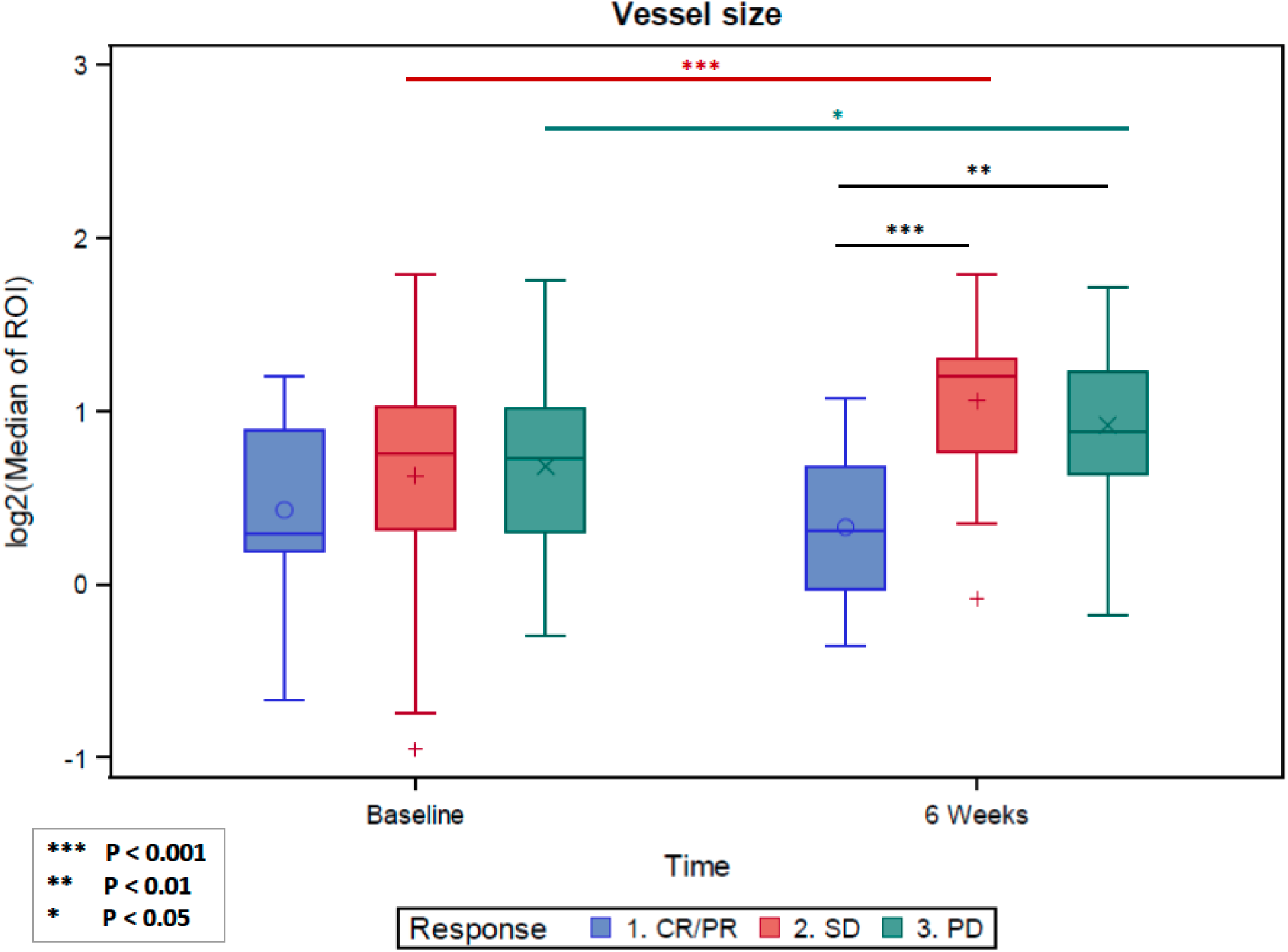
Pembrolizumab exerts differential effects on vessel size within BM depending on response. Blood vessels within ICI-responsive BM had smaller vessel sizes, compared to BM that did not respond to ICI. Median post-treatment vessel sizes were significantly larger for BM with SD (p=0.0004) and PD (p=0.002), compared with PR/CR. The median vessel size increased during treatment with pembrolizumab for patients with SD (p=0.0006) and PD (p=0.02). No change was noted for PR (p=0.43) in the same interval.

**Figure 4:**
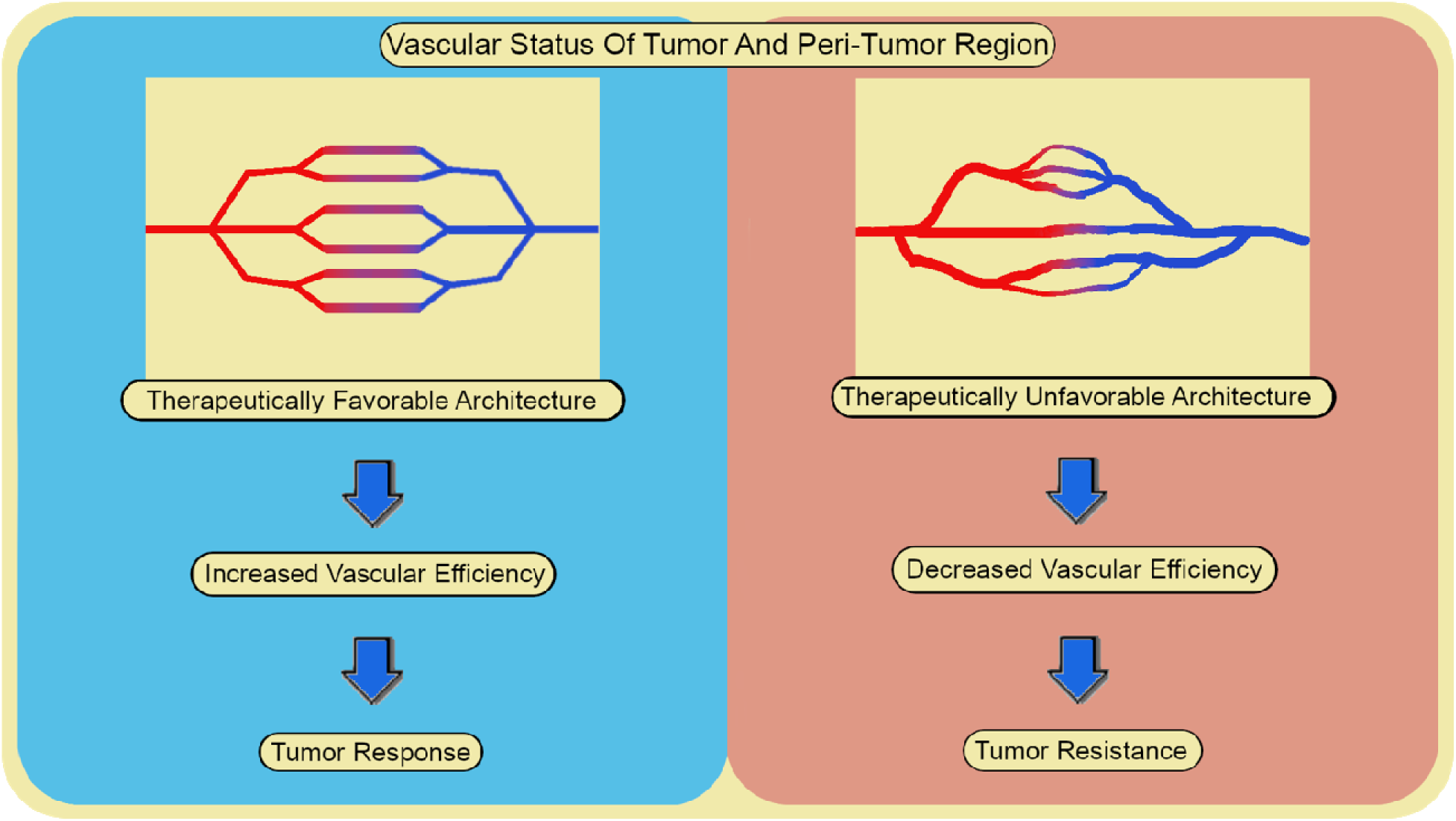
Schematic illustration of pre-treatment vasculature in ICI-responsive BM (left), compared with dysfunctional vasculature observed in ICI-resistant BM (right). A ‘therapeutically favorable’ vasculature harbors a more normal-functioning vascular architectur and stable blood flow, which will augment immune activity and the therapeutic impact of ICI. A ‘therapeutically unfavorable’ vasculature possesses abnormal vascular structures and increased vascular permeability, resulting in decreased vascular efficiency (e.g. vessel leakiness, unstabl blood flow, hypoxia, acidity) – thus reducing immune activity and the therapeutic impact of ICI.

We did not identify a significant relationship between microvascular CBV and treatment outcomes. This finding may be attributable to two factors: vessel distension, the presumed mechanism of increase in vessel size associated with ICI resistant tumors, and the low signal-to-noise ratio (SNR) for SE data as minimal contrast permeates the microvasculature. Furthermore, there was a trend (p=0.13) towards a longitudinal decrease in under-perfused tissue for ICI-responsive BM, suggesting that tumor hypoxia may be reversed in effective ICI (**Figure 2** – right panel; **Supplemental Figure 5**). Consistent with our finding that ICI-resistant BM possess a vascular architecture dominated by large-caliber vessels, PR/CR BM, compared to SD or PD BM, displayed a trend towards a potentially more therapeutically favorable level of under-perfused voxels (i.e. less under-perfused, hypoxic regions) at both pre-treatment and post-treatment (**Figure 2** – right panel). Next, we measured vascular permeability through the permeability parameter K^trans^, which has been linked to tumor grade and aggressivity in gliomas^39^. ICI-responsive BM had a significant increase in K^trans^ (p=0.02) during treatment (**Supplemental Figure 6**). No significant changes were seen in BM with SD or PD.

Finally, we obtained pre-pembrolizumab BM tissue for 12 patients enrolled in our study. We estimated immunogenicity and relative sizes of blood vessels, if present, using archival hematoxylin and eosin (H/E)-stained slides. Tissue from BM that derived intracranial benefit from pembrolizumab had a high density of tumor-infiltrating lymphocytes (TILs) and a mix of small and medium blood vessels (**Supplemental Figure 7**). Conversely, tissue from BM that were resistant to pembrolizumab had abundant tumor cells, few TILs, and a higher degree of large-caliber blood vessels (**Supplemental Figure 8**). Taken together, these histopathological and functional imaging findings suggest that vascular structural architecture is linked to immunogenicity and intracranial ICI outcomes.

### Unbalanced vascular architecture within the peri-tumor area is linked with ICI resistance

There was no difference in any pre-treatment vascular biomarker in the peri-tumor analysis for each response category. However, ICI-resistant BMs had highly significant increases over time (p<0.0001) for the fraction of microvasculature and under-perfused tissue (**Figure 5**). Similarly, there were non-significant decreases for these parameters towards a potentially more favorable level within ICI-responsive BMs. Furthermore, the rates of change for the level of microvasculature and under-perfused tissue from pre-treatment to post-treatment were highly different for PR/CR vs. SD/PD (p=0.008), with increases in PD BM compared to slight decreases or stability for the PR/CR BM, suggesting that an increase in ‘therapeutically unfavorable’ vasculature heralds tumor progression and lack of response to ICI (**Figure 4**).

**Figure 5:**
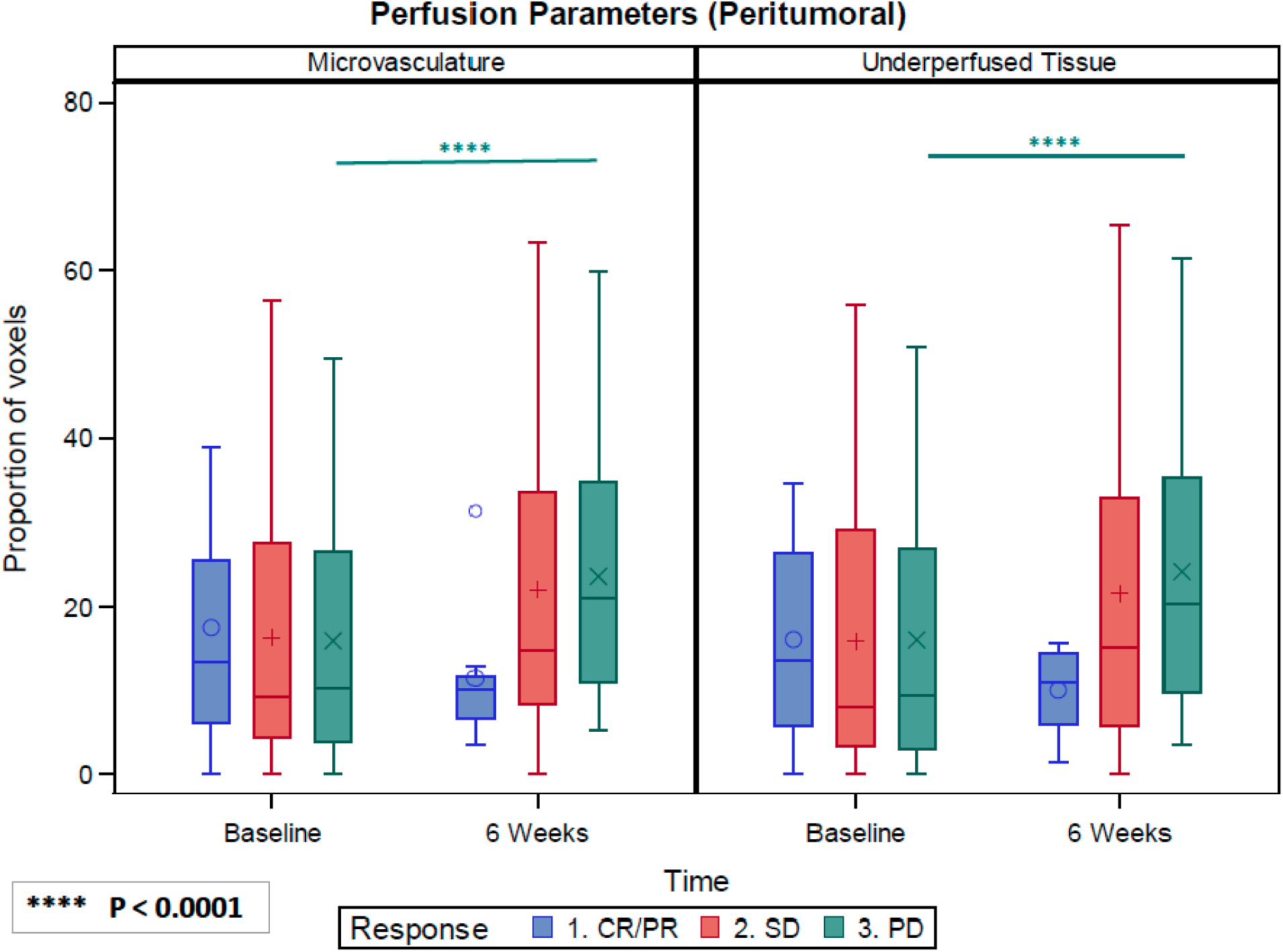
DSC-MRI peri-tumor analysis may detect early tumor progression before development of contrast enhancement. For both perfusion parameters, there were significant interactions between ICI response and visit. BM with PD had highly significant increases over time (p<0.0001) for both parameters, with no significant changes noted for PR or SD. Furthermore, the rates of change from baseline to 6 weeks were different for PR vs. PD (p=0.008 – microvasculature; p = 0.008 – under-perfused tissue).

### Arterial and venous physiology within BM

We found no difference in the fraction of arterial or venous-dominant voxels within BM between intracranial response and resistance, nor were there notable changes over time (**Supplemental Figure 9**). To assess for arterial or venous-specific physiologic changes that pembrolizumab might exert, we quantitated GE CBV/CBF and SE CBV/CBF for the arterial or venous-dominant voxels within the tumor. There were no significant trends for any parameter with respect to intracranial response (**Supplemental Figure 10**).

### Assessment of inter-patient and inter-histology variability in vascular physiology

We assessed heterogeneity in vascular parameters within multiple BM of the same patient and focused our analysis on patients that had at least 3 or more BM assessed in this study. Vascular biomarkers for BM of the same patient were more similar to each other, compared to those for BM of different patients (**Supplemental Table 2**). Next, we assessed histology-specific differences in vascular physiology. No significant difference in any vascular biomarker, according to histology, was identified (**Supplemental Table 3**).

### Non-vascular multi-parametric assessments of BM

We also performed analyses on fractional anisotropy (FA) and apparent diffusion coefficient (ADC) on all BM with evaluable data. We found no difference in median FA within the contrast-enhancing and peri-tumor regions between BM with intracranial response versus resistance, nor were there notable changes over time (**Supplemental Figure 11**). There was a statistically significant increase in median ADC for both the contrast-enhancing region (p = 0.04) and peri-tumor region (p = 0.006) of PD BM between the pre-treatment and post-treatment visit (**Supplemental Figure 12**). No significant differences pre-treatment and post-treatment were observed for BM with SD or PR/CR.

## Discussion

We first introduced the concept of a ‘therapeutically unfavorable’ vascular architecture mediating radiation necrosis in prior work^24^, and now extend this paradigm to immunomodulation and ICI efficacy. Here, we found that that a structurally balanced and functional vascular network may contribute to ICI response. On radiographic and histological analysis, ICI-responsive BM possessed a balanced mix of both small and large-caliber blood vessels and an inflammatory TME. Conversely, ICI-resistant BM possessed an immunosuppressive TME dominated by large-caliber vessels, consistent with prior studies showing increased vessel size in advanced cancers^15–17^. Moreover, we identified a longitudinal trend towards a decrease in under-perfused tissue in BM responding to ICI, suggesting a reversal of the negative effects of hypoxia^15^. Taken together, responding tumors may display a shift towards a favorable balance of small and large-caliber blood vessels that promote vascular efficiency^24,40^ and therapeutic immune modulation. By contrast, a TME dominated by large-caliber vessels may represent an advanced phase in tumor evolution and not be a favorable environment to maintain sufficient vascular connectivity for treatment response^30,41,42^.

These results are consistent with tissue-based and functional imaging studies for brain tumors that illustrate a link between abnormal vasculature and aggressive phenotypes. Histopathological analysis of high-grade gliomas and BM has demonstrated thick, dilated, and glomeruloid microvasculature, compared to lower-grade tumors or normal brain tissue^22,43^. Additionally, an orthotopic murine model revealed that glioblastoma progression is associated with a dramatic increase in vessel diameters and accumulation of immunosuppressive macrophages^44^. Another study in glioblastoma patients treated with bevacizumab found that long-term survivors exhibited an overall decrease in vessel caliber^22^. By contrast, rapid progressors had a higher degree of larger-caliber venule-like vessels^22,42^. Our study, which is one of the largest DSC-MRI studies performed in human BM, provides further evidence linking a balanced vascular architecture to increased vascular efficiency and a pro-inflammatory, anti-tumor TME that augments ICI efficacy. While our study defined an estimate for a ‘therapeutically favorable’ vascular architecture for ICI, we note that a ‘favorable’ balance of small- and large-caliber vessels may vary depending on vascular characteristics of reference tissue, tumor histology, and prior therapy^30,45,46^. Future studies in more homogenous cohorts may delineate exact thresholds for ‘favorable’ and ‘unfavorable’ vascular architectures for specific patient populations.

In contrast to prior studies suggesting decreasing vascular permeability associated with treatment response^40,47^, we found an increase in permeability in ICI-responsive BM. This phenomenon may be due to a local inflammatory response exerted by pembrolizumab that may result in increased permeability and immune cell efflux into BM. Similarly, another recent study demonstrated increased vascular permeability in glioblastoma patients who responded to bavituximab, an immune-activating agent^48^. The beneficial increase in permeability seen with bavituximab and pembrolizumab may enhance CNS penetration of these large molecular weight antibodies that may otherwise have difficulty in crossing the blood-brain barrier.

Importantly, perfusion-weighted imaging enables investigation of the peri-tumor region, which is generally less studied than the tumor core. In the peri-tumor region of ICI-resistant BM, the percentage of small vessels and hypoperfusion increased following treatment. These findings were identified prior to development of contrast enhancement, suggesting neo-angiogenesis from micro-metastatic tumor progression and a shift toward an unbalanced and less favorable vascular network as the tumor starts to grow. Consistent with studies in other tumor types^49^, the peri-tumor area of BM has divergent biology compared to that of contrast-enhancing tumor, suggesting that there is germane tumor biology to be gleaned from the peri-tumor area. An increased fraction of microvasculature within contrast enhancement may indicate a shift towards fewer established abnormally dilated vessels and therefore a more ‘therapeutically favorable’ vascular architecture. Conversely, an increase in microvasculature within the peri-tumor area may herald neo-angiogenesis and burgeoning tumor growth.

Our study had several limitations. First, the FOV for the DSC sequence was limited and encompassed only slightly more than half of the brain parenchyma in order to optimize temporal and spatial resolution^50^. In addition, because of the limited size of the FOV, a BM captured on one visit was not always captured on future visits. This restricted the number of BM for which we had longitudinal evaluable data. After our study was planned, a new DSC GE/SE sequence was developed that enables whole brain coverage for high-resolution VAI; using this sequence should be considered for future studies^50^. Furthermore, as ICI received approval for melanoma and NSCLC BM soon after our trial started^17,18^, we had a heterogenous dataset that encompassed 11 different histologies who could not receive ICI as part of clinical care. While we did not detect overt differences in vascular biomarkers according to histology, this heterogeneity may preclude discovery of subtle differences. Prospective validation studies in well-defined cohorts (e.g. homogenous cohort of histologies, pre-SRS vs post-SRS) are needed.

While we found co-existence of increased TILs and a structurally balanced vascular architecture, we were not able to obtain more detailed insights of immune cell function. Given the challenges in sampling spatially separated BM serially, future efforts optimizing these non-invasive techniques in probing germane facets of the TME are critical. Additionally, as this was a single-arm study, we do not know whether identified patterns in vessel architecture are specific for ICI efficacy, or simply reflective of TME facets generally seen in tumor response or resistance. Further work is needed before employing vascular architecture as a therapy agnostic tool for assessing tumor biology. Finally, we note that vascular physiology is only one facet of the TME that contributes to ICI efficacy. Future gene expression profiling efforts for immune cell populations (e.g. effector T-cells, myeloid-derived suppressor cells) may help determine molecular or transcriptional factors that link specific patterns of vessel architecture with immune function. Strategies that combine immunomodulation with agents that shift a dysfunctional vascular architecture (e.g. bevacizumab) to a normalized and therapeutically favorable network may hold promise.

In summary, we performed pre- and post-treatment physiologic imaging to investigate how structural and functional vascular physiology contribute to tumor immunogenicity and ICI efficacy in BM. Our results suggest that certain vascular architectures improve vascular efficiency, which is linked with improved immune function and ICI efficacy. While these findings have promise for shedding light on mechanisms of response, they require additional validation. Future studies combining Omics-based and functional studies that illuminate the mechanisms linking patterns of vascular architecture with immunomodulation may inspire future combination treatment strategies.

## Methods

### Study design

All patients were enrolled on a phase 2 trial (NCT02886585) evaluating intracranial efficacy of pembrolizumab. Participants had measurable intracranial disease, defined as at least one lesion that measured in at least one dimension as > 5 mm, and histologically confirmed disease from any solid tumor. Complete eligibility criteria are noted in [companion manuscript citation]. All patients were treated with 200 mg of intravenous pembrolizumab every three weeks, underwent a baseline MRI examination, and follow-up MRI’s every 6-8 weeks (**Figure 1**).

### MRI acquisition and tumor segmentation

All MRI examinations were performed on a 3T Skyra (Siemens Healthineers, Erlangen, Germany). The MRI protocol included 3-dimensional T1-weighted MPRAGE images pre- and post-contrast injection, T2-weighted fluid attenuated inversion recovery (FLAIR), diffusion weighted imaging, dual-echo DSC MRI with simultaneous gradient echo (GE) and spin echo (SE) acquisitions, and dynamic contrast enhanced (DCE) imaging. All sequences were acquired in the same session. The dual-echo spiral DSC sequence was acquired with the following parameters: TR/TE1/TE2 = 1500/31/104 ms; flip angle = 60 degrees; field of view (FOV) = 240-264 x 180-198 mm, slice gap = 6.5 mm, number of slices = 11, and acquired matrix = 120 x 90.

Structural images (FLAIR, pre-/post-contrast MPRAGE) and parameter maps from DTI, DSC, and DCE imaging were registered to the T2-weighted image using the BRAINSFit module in 3D Slicer^51^. Contrast enhancement, excluding central necrosis, was delineated on the post-contrast MPRAGE images by using DeepMETS, a deep learning algorithm that automatically segments contrast-enhancing brain tumors^52^. These regions-of-interest (ROIs) were reviewed and manually adjusted as needed (A.E.K. & E.R.G.). Each BM were manually annotated and tracked longitudinally across visits. BM were assessed for intracranial response using Response Assessment for Neuro-Oncology for Brain Metastases (RANO-BM) criteria^53^. The peri-tumor region-of-interest (ROI) was created by a 4-mm wide dilation of the tumor ROI and excluding the original central, contrast-enhancing, region. This margin to define the peri-tumor region was chosen to encompass a biologically relevant area around the tumor as noted in a prior study^24^.

### Image processing

NordicIce (NordicNeuroLabs AS, Bergen, Norway) and VAI were used to generate leakage-corrected GE and SE cerebral blood volume (CBV), cerebral blood flow (CBF), vessel size, and arterial or venous-dominant voxel maps (**Table 1**)^22,24,36^. SE-derived maps reflect microvasculature, while GE-derived maps represent pan-vascular or both micro- and macrovascular perfusion^54^. Arterial or venous-dominant voxel maps are generated from a temporal shift between the contrast agent induced signal change of the DSC GE and SE images^22^. This shift can be quantified by a point-by-point parametric plot to form a vortex, were the direction of the vortex depended on the type of vessels included in the vascular system. In a five-level split, voxels with the largest positive shift (clockwise direction) are associated with arterial dominated regions, and voxel with the largest negative shift (counter-clockwise direction) is associated with venous dominated regions. Consistent with prior work^24,38,42,55–57^, all MRI-derived maps were normalized to reference tissue using NordicIce. As BM encompass both grey matter (GM) and white matter (WM), reference tissue was defined by the WM and GM masks^22,30,42,58–62^, excluding areas with contrast enhancement and abnormal T2-weighted/FLAIR hyperintensity. All analyses were performed with normalized values^38,55–57,63^, which is a unit-less ratio of absolute voxel values to the reference tissue. DTI data were processed to generate fractional anisotropy (FA), apparent diffusion coefficient (ADC), values and corresponding ADC maps, using DTIFit from the FSL Diffusion Toolbox. As the log scale modeling provided better assessments of fit due to Akike information criterion and simulation-based calibration, all figures were presented on the log scale. This technique more accurately illustrates the variability of the data within the models.

Given the link between angiogenesis and immunosuppression, we specifically studied structural and functional aspects of tumor microvasculature. Per prior work^24^, we assessed the fraction of the total tumor vasculature and perfusion below the 20^th^ (<20%) percentile of the average pre-treatment normal appearing tissue across all patients. This threshold is a good indicator of the level of microvasculature (pan-vascular [GE] CBV <20%)^64,65^ and under-perfusion (pan-vascular CBF <20%) within the tissue (**Table 2**). These analyses enable a better characterization of tumor heterogeneity for the CBV/CBF parameters, rather than simply the median.

**Table 2:** Biological interpretation of vascular parameters.

| **Vascular Parameter** | **Biological meaning** |
| --- | --- |
| GE CBV | Aggregate cerebral blood volume within tumor across all vessels |
| GE CBF | Aggregate cerebral blood flow within tumor across all vessels |
| SE CBV | Aggregate cerebral blood volume, specific for microvasculature |
| SE CBF | Aggregate cerebral blood flow, specific for microvasculature |
| Microvasculature | Voxels characterized by small-caliber vessels (GE CBV under the 20<sup>th</sup> percentile of average pre-treatment normal appearing tissue across all patients) |
| Under-perfused tissue | Voxels characterized by hypoperfusion (GE CBF under the 20<sup>th</sup> percentile of average pre-treatment normal appearing tissue across all patients) |
| Vessel size index | Mean vessel diameter (or caliber) within a voxel of interest |
| Arterial-dominant voxels | Voxels in which arteries are the dominant vessel type (voxels with positive vortex direction) |
| Venous-dominant voxels | Voxels in which veins are the dominant vessel type (voxels with negative vortex direction) |
| $K^{trans}$ | Capillary permeability – measured by the accumulation of contrast in the extravascular-extracellular space |

In addition, we deferred a SE-specific assessment for the <20% analyses given the inherent low signal-to-noise ratio (SNR) of the SE signal in such small vessels^54^. To define microvasculature, we used the bottom 20^th^ percentile of GE CBV, rather than SE CBV, to maximize the robustness of our exploratory findings, given the relatively small sample size and low SNR of SE CBV. A minimum of 10 non-zero voxels within the ROI was established as a threshold to be included for analysis.

### Histopathological assessment

We identified archival BM tissue resected from enrolled patients prior to treatment with pembrolizumab. A board-certified pathologist confirmed the histologic diagnosis and selected representative formalin-fixed paraffin-embedded samples with high tumor purity. Using a hematoxylin-and-eosin stained slide, the pathologist qualitatively estimated tumor-infiltrating lymphocyte (TIL) burden and, if present, relative sizes of blood vessels.

### Statistical analysis

For statistical analysis of all vascular physiology metrics, apparent diffusion coefficient (ADC), and fractional anisotropy (FA), the unit of analysis is the individual BM. Analyses were performed on the level of the BM, and not the patient, given the incidence of mixed intracranial response between spatially-separated BM in the same patient (e.g., where 1 BM will respond but others will progress; existing BM respond but a new BM develops). Since patients could have more than one BM, the analysis allows for clustering of BM within patient. To investigate the relationships with metastasis-level response by visit and over time, longitudinal linear mixed models were fit with log2(median measurement) [or log2(percentage) for arteries and veins] as the dependent variable, and visit, response, and their interaction as independent predictors. If the outcome value was zero, one was added prior to taking the log2 transform. The models used compound symmetry covariance to estimate the within-BM relationships over time. Pairwise differences between response categories (PR/CR, SD, or PD) as well as changes over time were estimated using contrasts. Changes are expressed as log2(fold-change).

To assess relationships between vascular biomarkers and histology, patients were classified according to histology: breast, lung, and other. These models expanded the original longitudinal modeling and regressed log2(median) on main effects of histology, response, visit, and the interaction of visit and response. The p-values reported are for the main effect of histology controlling for the other factors. As in the previous modeling, the BM was the unit of analysis, and the data were clustered by patient to allow for within-patient correlations using compound symmetry.

To assess heterogeneity of vascular biomarkers for BM within the same patient, we focused our analysis on the 17 patients with 3 or more BM used for this analysis. We used the individual BM mean and ROI data to calculate the coefficient of variation (CV; defined as STD/mean) for each BM <u>at baseline</u>. This is a unitless measure and shows the extent of variability relative to each mean. CV is preferred rather than the STD alone because variability can increase as the mean increases. Separate repeated measures models were fit regressing log2(CV) upon patient or histology to assess for differences in mean CV. Multiple pairwise comparisons were adjusted using Tukey-Kramer. The p-value assessed if there were differences in average CV between patients or histology.

All analyses were performed using SAS 9.4 (SAS Institute Inc., Cary, NC, USA).

---

**Supplemental Figure 1:**
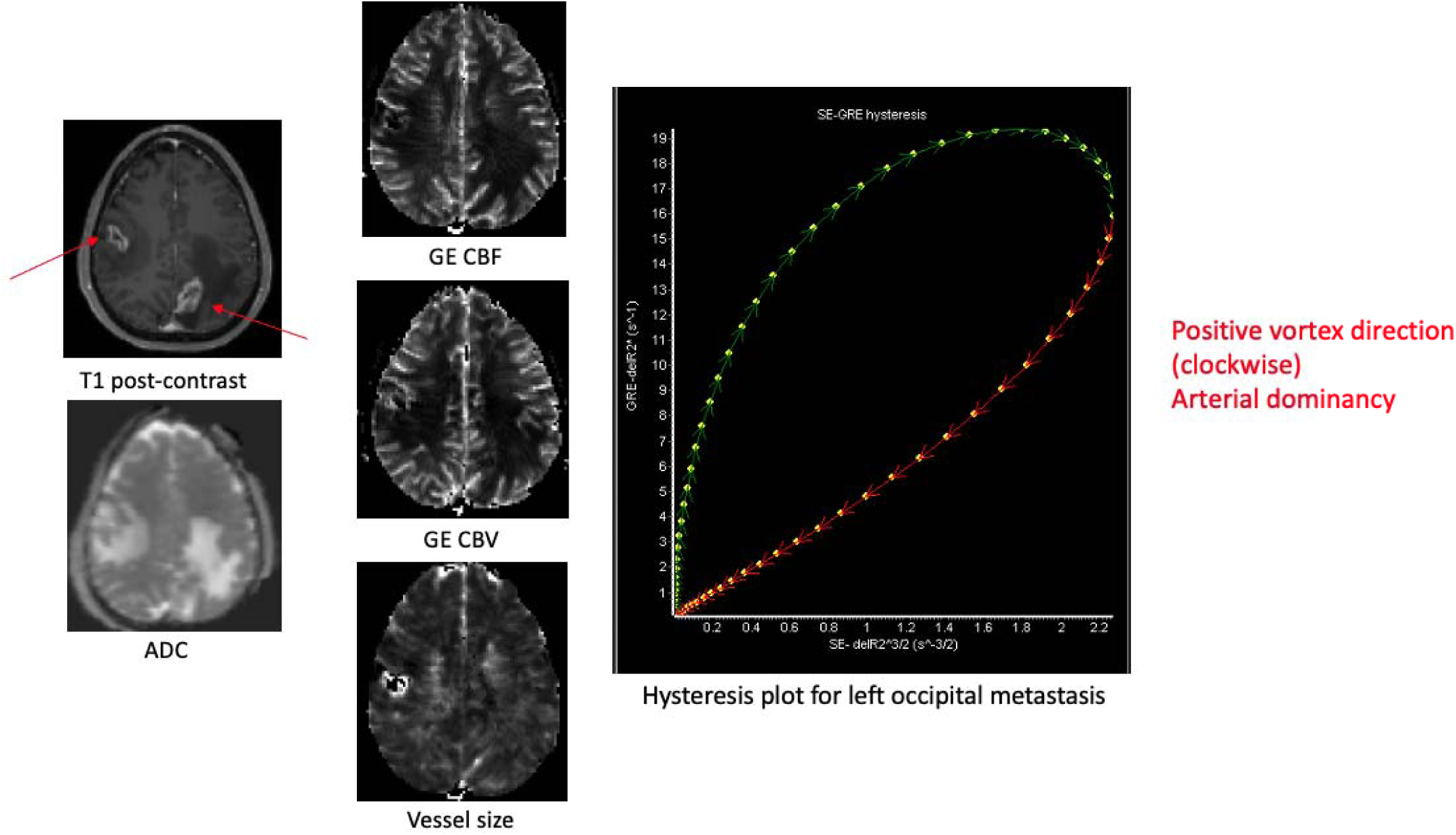
Parametric images and hysteresis plot from an occipital non-small cell lung cancer BM that responded to pembrolizumab. The fitted curves on the hysteresis plot are plotted with delta R2^3/2 (SE-based) along the x-axis and delta R2* (GE-based) along the y-axis. The clockwise rotation indicates that GE-based changes precede SE-based changes, corresponding to a negative peak shift and arterial dominance.

**Supplemental Figure 2:**
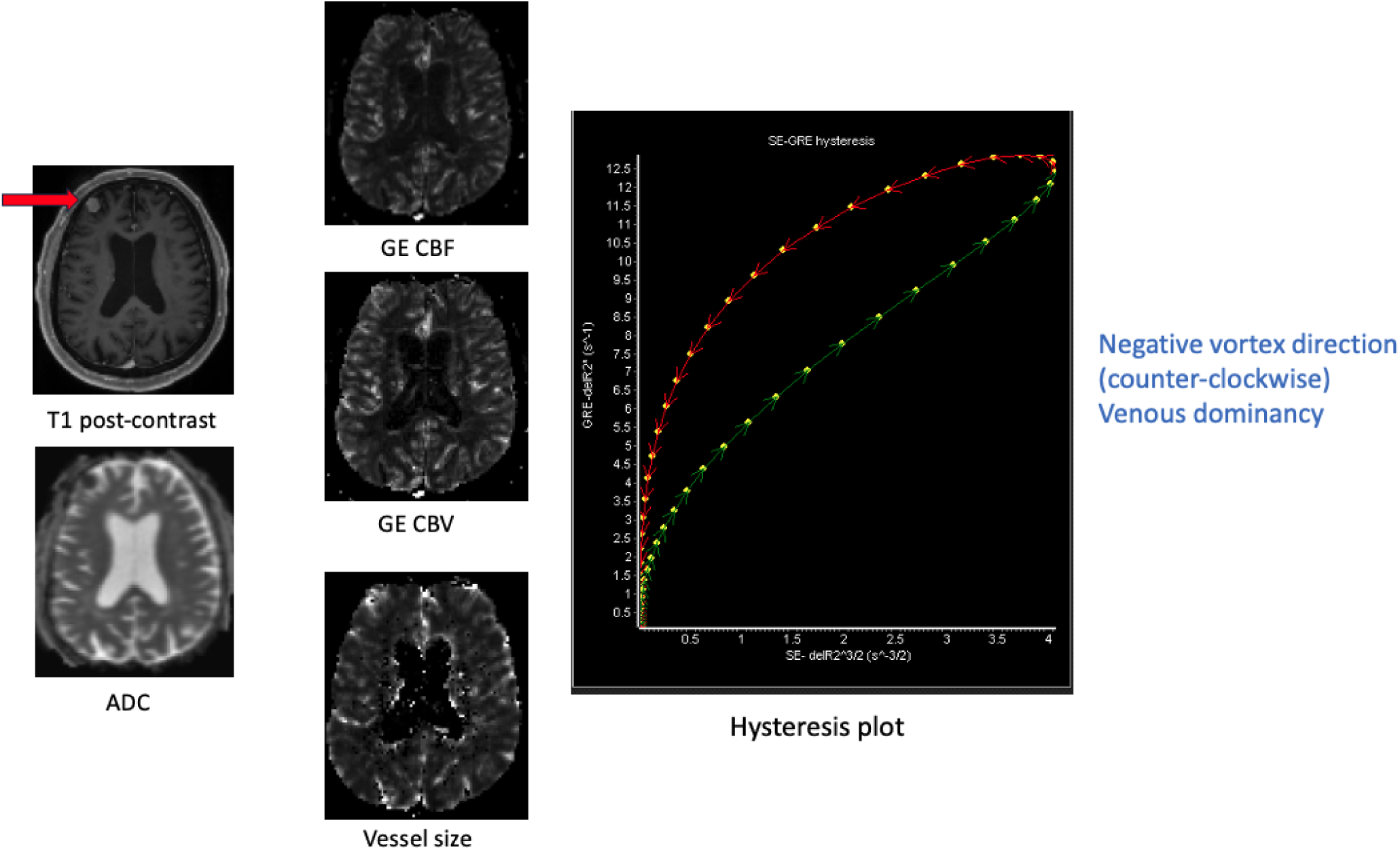
Parametric images and hysteresis plot from a frontal small-cell lung cancer BM with prolonged SD to pembrolizumab. The fitted curves on the hysteresi plot are plotted with delta R2^3/2 (SE-based) along the x-axis and delta R2* (GE-based) along the y-axis. The counter-clockwise rotation indicates that SE-based changes precede GE-based changes, corresponding to a negative peak shift and venous dominance.

**Supplemental Figure 3:**
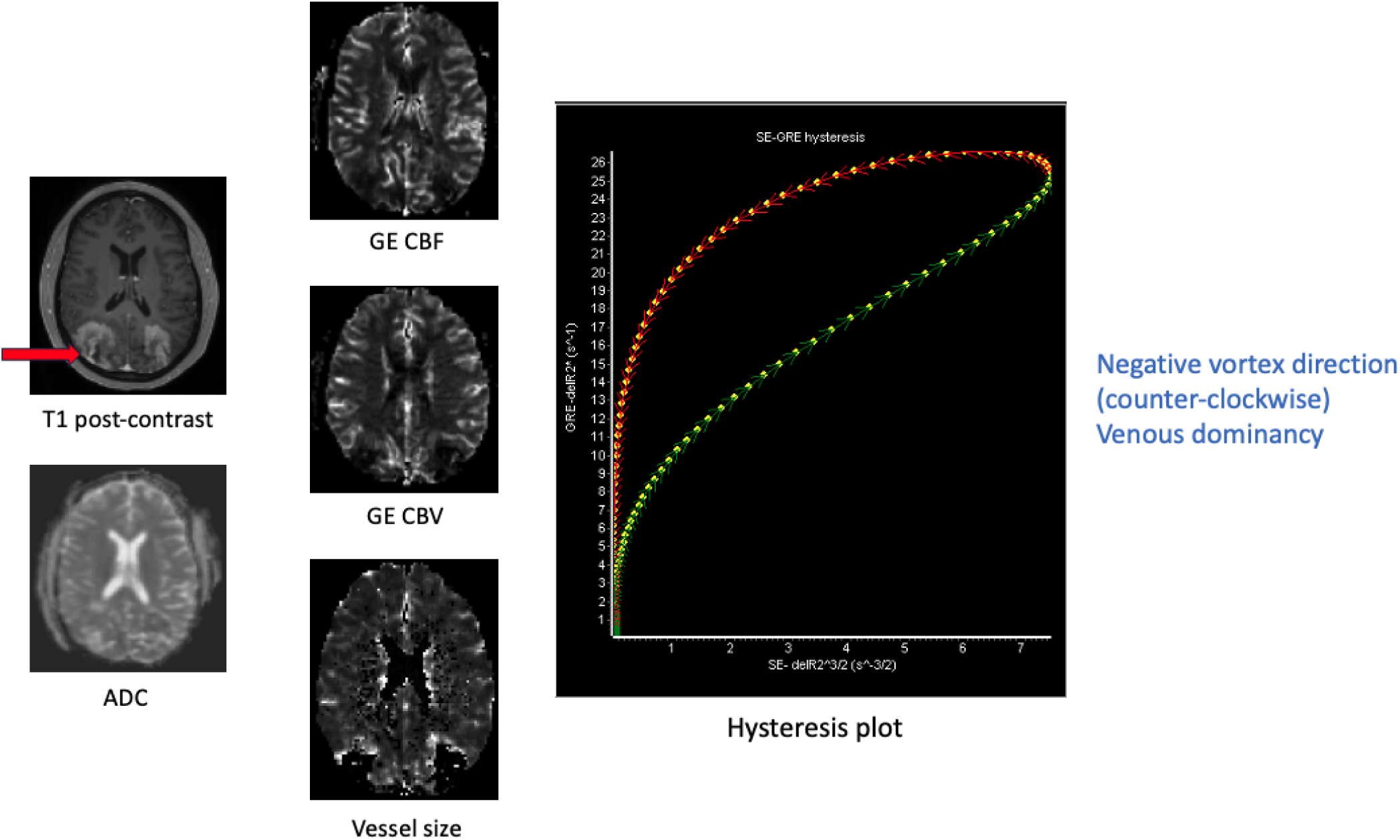
Parametric images and hysteresis plot from an occipital breast cancer BM with immediate PD to pembrolizumab. The fitted curves on the hysteresis plot ar plotted with delta R2^3/2 (SE-based) along the x-axis and delta R2* (GE-based) along the y-axis. The counter-clockwise rotation indicates that SE-based changes precede GE-based changes, corresponding to a negative peak shift and venous dominance.

**Supplemental Figure 4:**
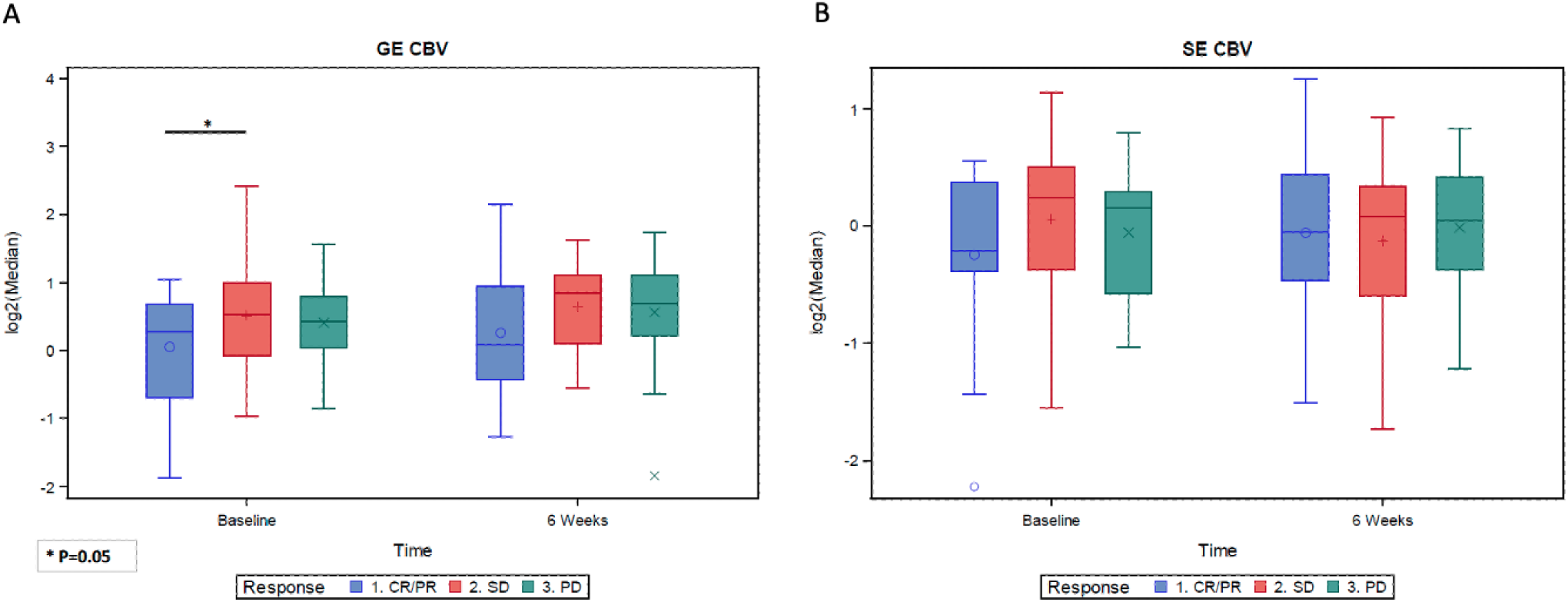
Median gradient-echo (A) and spin-echo (B) cerebral blood volume within BM during treatment with pembrolizumab, organized by response. ICI-responsive BM had significantly lower median baseline GE CBV values compared with SD BM (p = 0.05). ICI-responsive BM had a non-significant trend towards a lower median GE CBV at baseline than PD BM (p = 0.07). There were no significant changes between pre-treatment and post-treatment for GE CBV, nor were the rates of changes different between response groups. There was no significant relationship between median SE CBV and response.

**Supplemental Figure 5:**
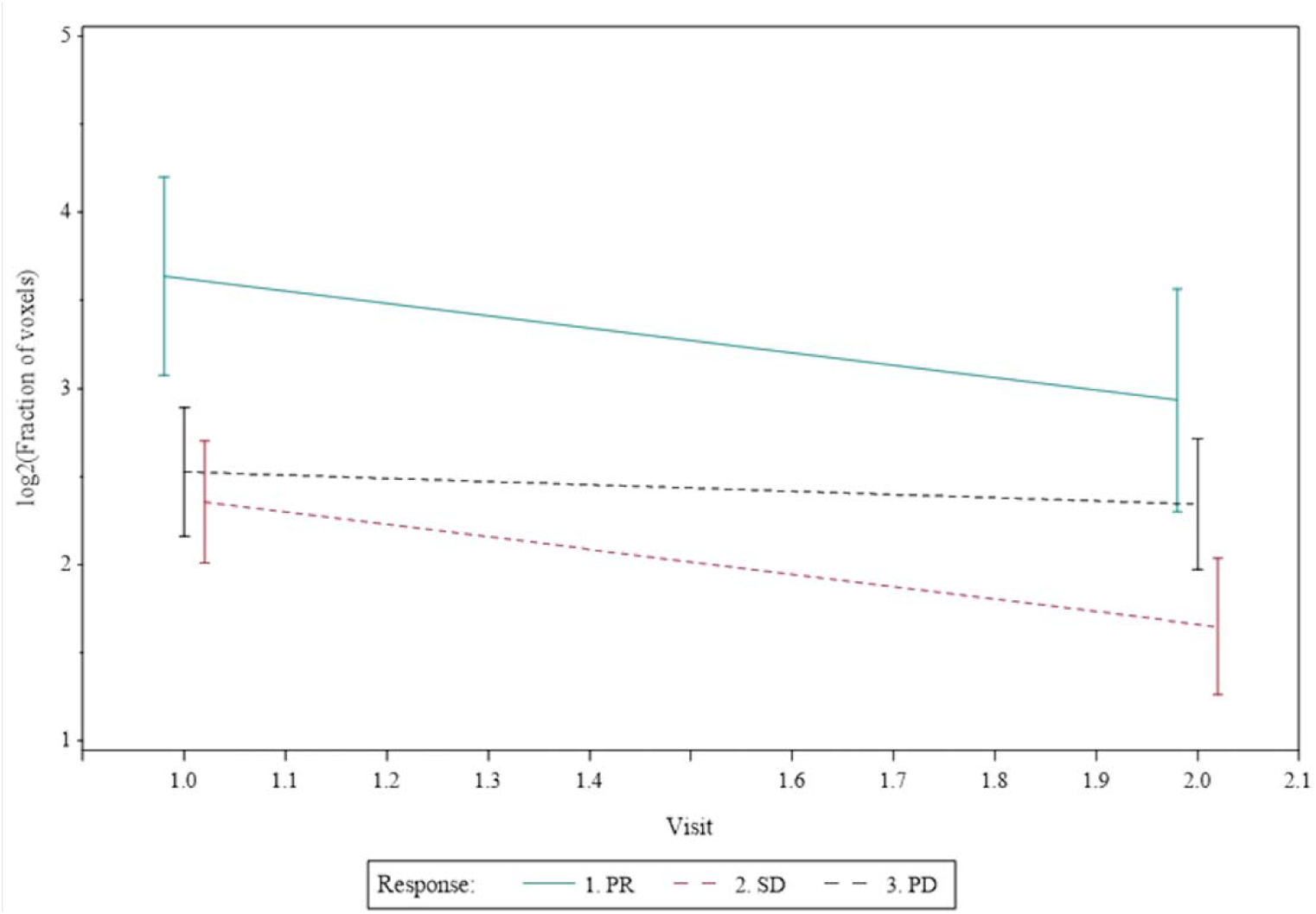
Assessment of under-perfused tissue within BM during treatment with pembrolizumab. This plot reflects the results of the longitudinal linear mixed models. The p-value for the longitudinal change in the CR/PR group (green line) is 0.13 and demonstrates the trend towards a decrease over time.

**Supplemental Figure 6:**
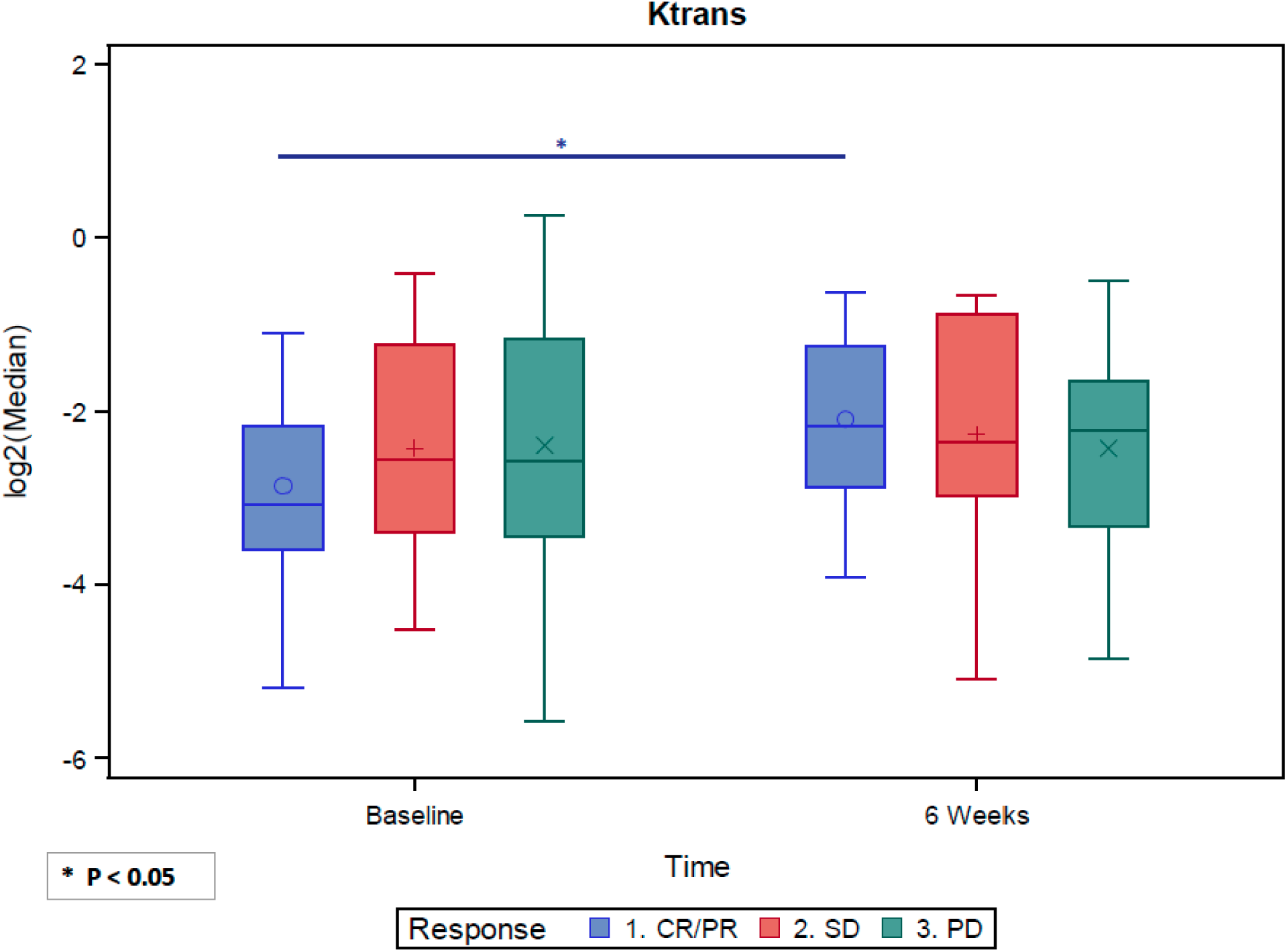
BM that responded to ICI are associated with an increase in vascular permeability. BM that responded to pembrolizumab had a statistically significant increase in K^trans^ between pre-treatment and post-treatment (p=0.02). This was not seen in BM with SD (p=0.58) or PD (p=0.75). While the analysis suggests that the longitudinal arc of change across visits between PR and PD are different, this did not achieve statistical significance (p=0.08).

**Supplemental Figure 7:**
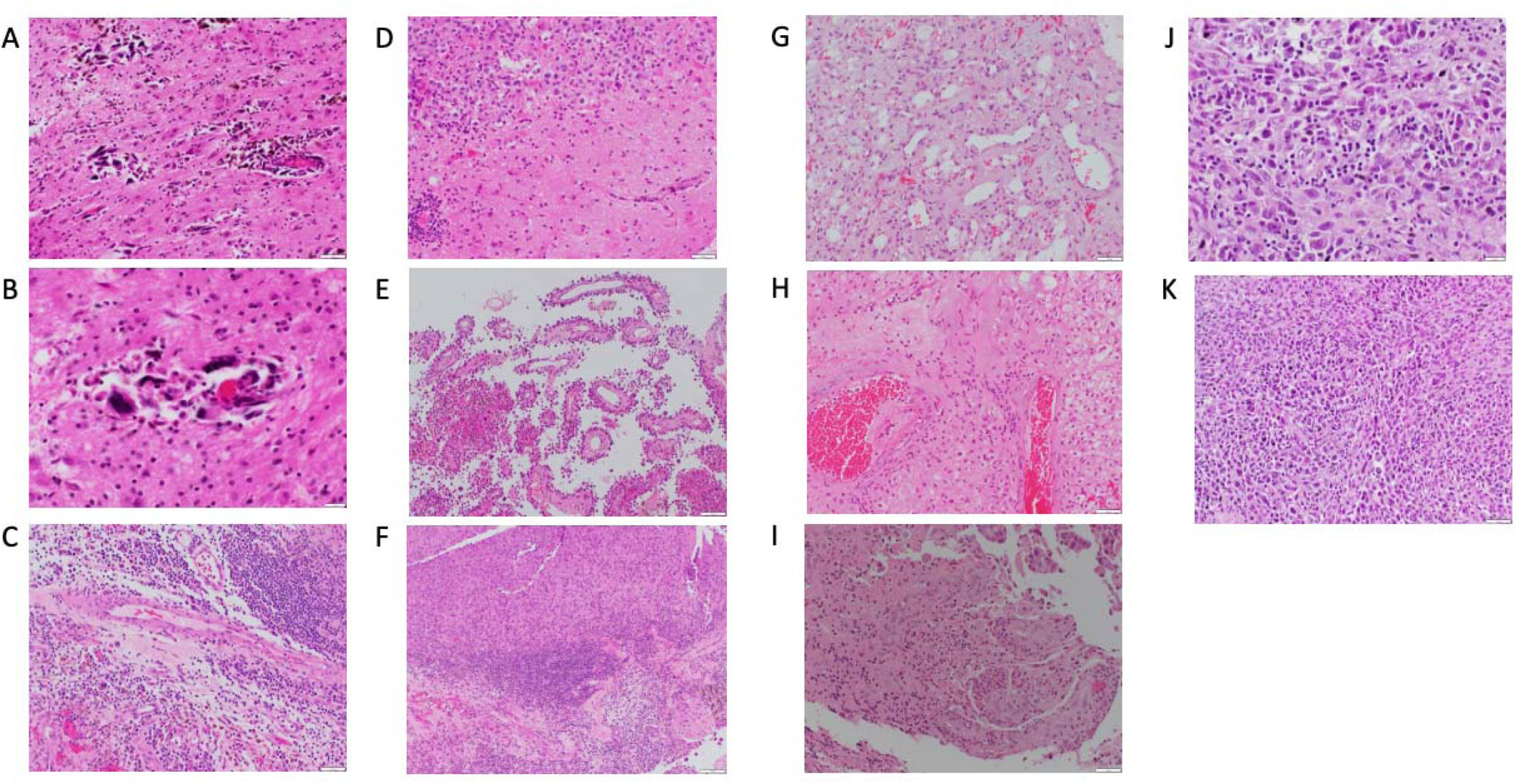
Representative H&E images from resected brain metastases that either had radiographic response or prolonged stability to pembrolizumab. A) and B) These samples are from a vermian BM (melanoma primary) and exhibit moderate TILs with relatively small caliber blood vessels in the periphery of the tumor surrounded by tumor cells. C) and D) These samples are from a left frontal BM (melanoma primary) and exhibits abundant TILs (C) with a normal-caliber blood vessel (C, middle) and smaller caliber vessels (C, bottom left). E) and F) These samples are from a BM (melanoma primary) and show tumor cells around small/medium caliber blood vessels with rare TILs (E), and in a different region many TILs including a lymphoid aggregate mixed with tumor cells and reactive changes (F). G) and H) These samples are from a left occipital BM (renal cell carcinoma primary), and exhibit an area of ectatic irregular vascular channels and surrounding reactive changes (G), and larger vessels with moderate perivascular TILs and macrophages, and occasional tumor cells. I) This sample is from a right parietal BM (breast primary) and exhibits tumor cells mixed with scattered TIL’s and neutrophils. J) and K) These samples are from a right cerebellar BM (rectal primary) and exhibit sheets of tumor cells and a moderate amount of TILs.

**Supplemental Figure 8:**
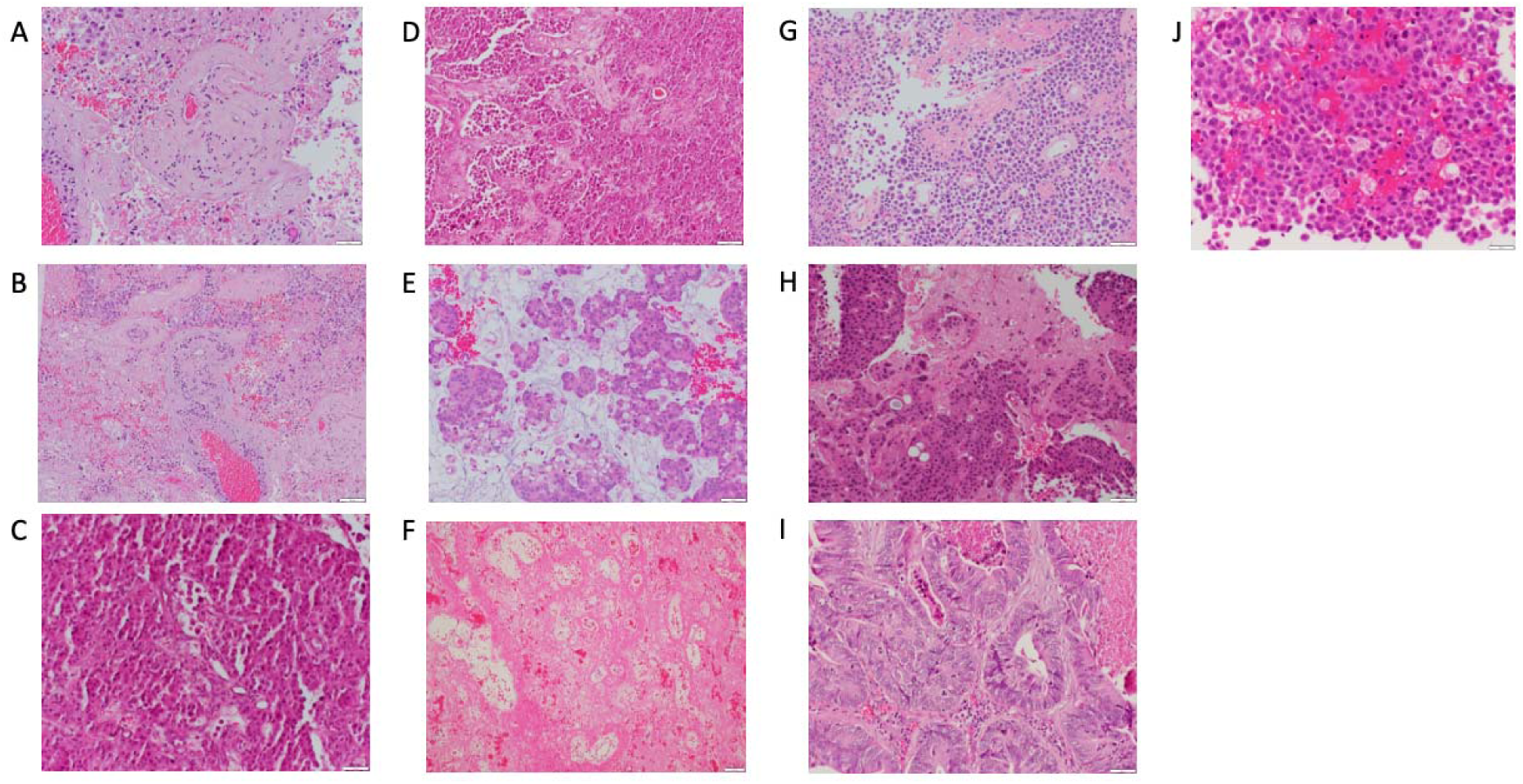
Representative H&E images from resected brain metastases that later were resistant to pembrolizumab on trial. A) and B) These samples are from a right parietal BM (breast primary) and exhibit many medium to large-caliber blood vessels with thickened walls, viable tumor with necrosis, and few TILs. C) and D) These samples are from a left cerebellar BM (breast primary) and demonstrate abundant tumor cells with minimal TILs. E) This sample is from a right occipital BM (esophageal primary) and exhibits an abundance of tumor cells with surrounding mucin and no inflammatory infiltrate. F) and G) These samples are from a left frontal BM (breast primary) and exhibit a diffuse fibrinous background with abnormal vasculature (F) and abundant tumor cells with focal interstitial fibrin and minimal TILs (G). H) This sample is from a right frontal BM (breast primary) and exhibit abundant tumor cells with minimal TILs and surrounding brain parenchyma. I) This sample is from a right frontal BM (colorectal primary) and shows abundant viable tumor with sparse TILs. J) This sample is from a right parietal BM (breast primary) and demonstrates abundant tumor cells with minimal inflammatory infiltrate.

**Supplemental Figure 9:**
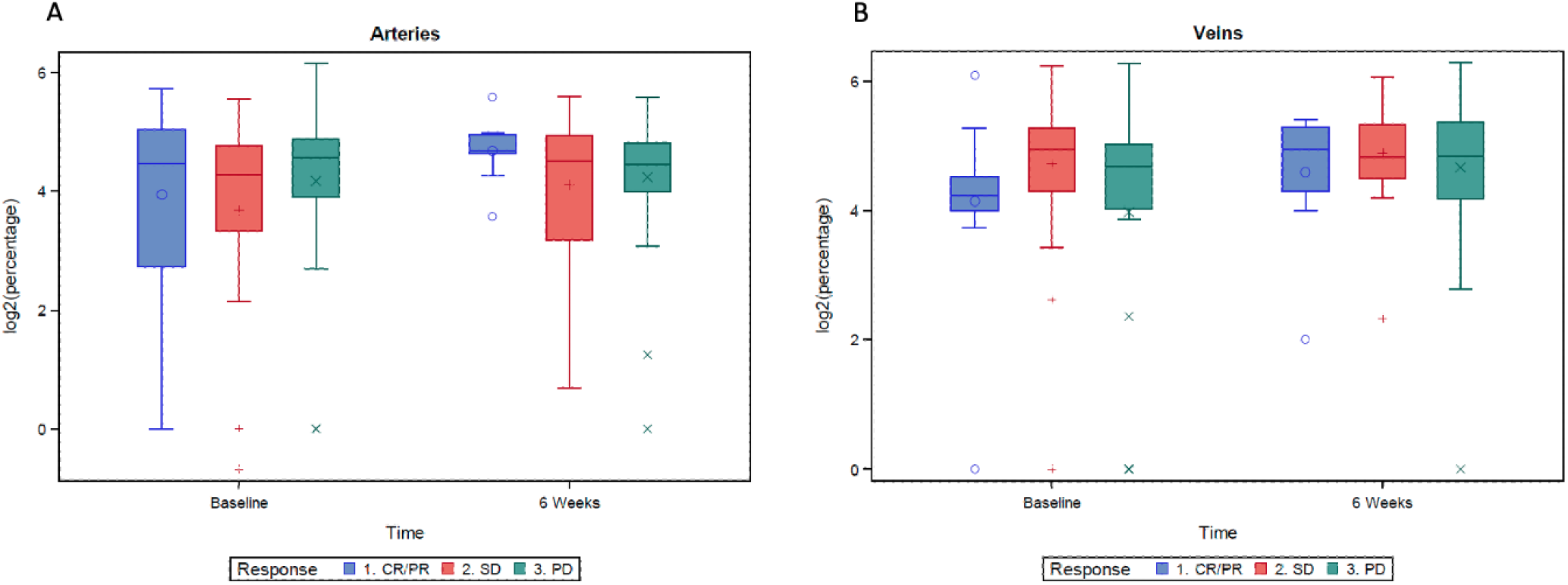
Arterial-dominated (A) and venous-dominated (B) voxel density within BM during treatment with pembrolizumab, organized by response. There were no statistical differences between responses in percentage of arteries and veins at either pre-treatment or post-treatment. BM that progressed on treatment had a non-significant increase in percentage of venous voxels between pre-treatment and post-treatment (p=0.06).

**Supplemental Figure 10:**
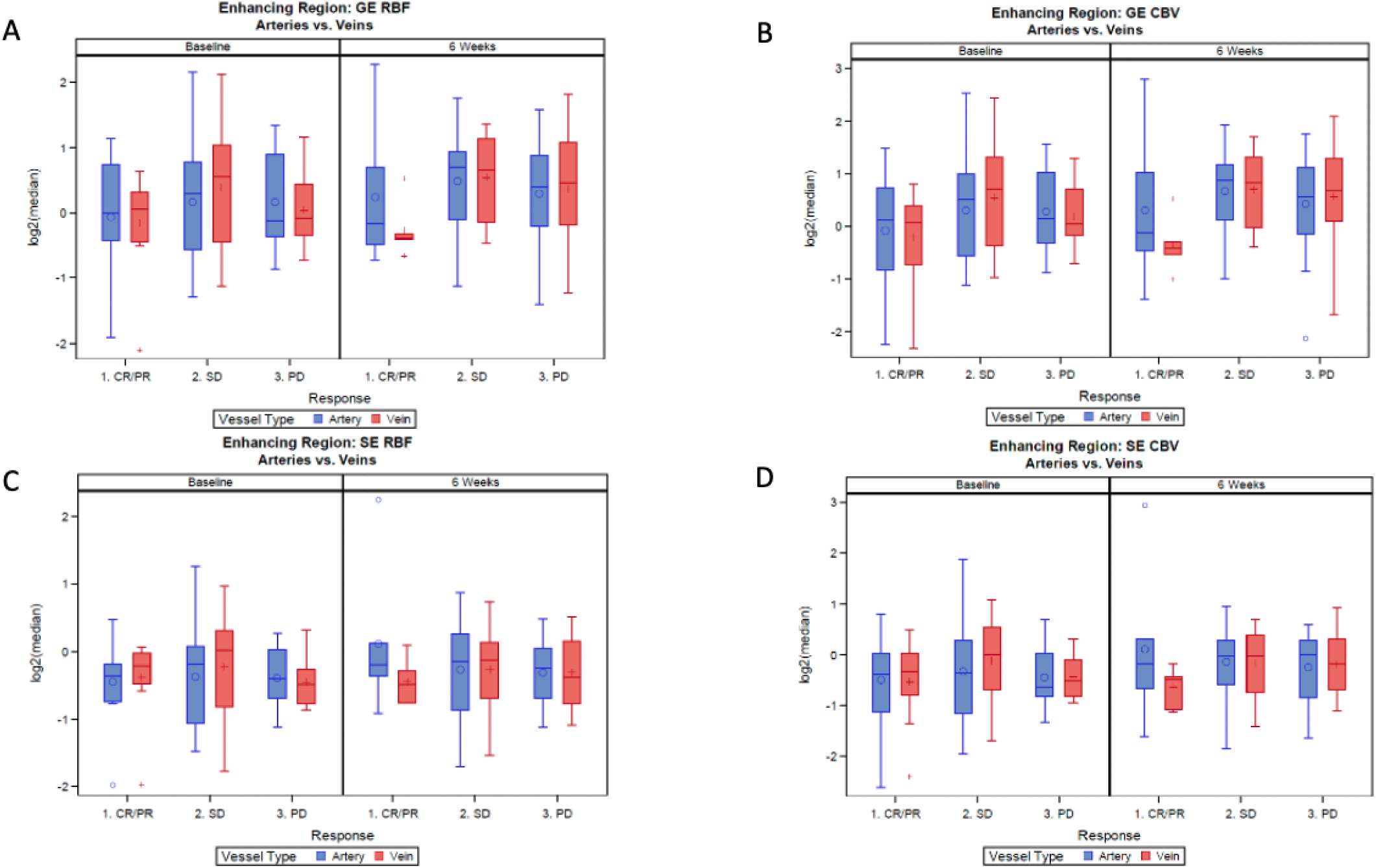
Vascular physiology within arterial and venous-dominated voxels in BM during treatment with pembrolizumab. Analyses to determine significant correlations in vascular physiology within arterial or venous-dominated voxels were performed. After Benjamini-Hochberg correction for false discovery rate, there were no statistically significant trends for any vascular parameter (GE CBF [A], GE CBV [B], SE CBF [C], and SE CBV [D]) within arterial- or venous-dominated voxels for each response category.

**Supplemental Figure 11:**
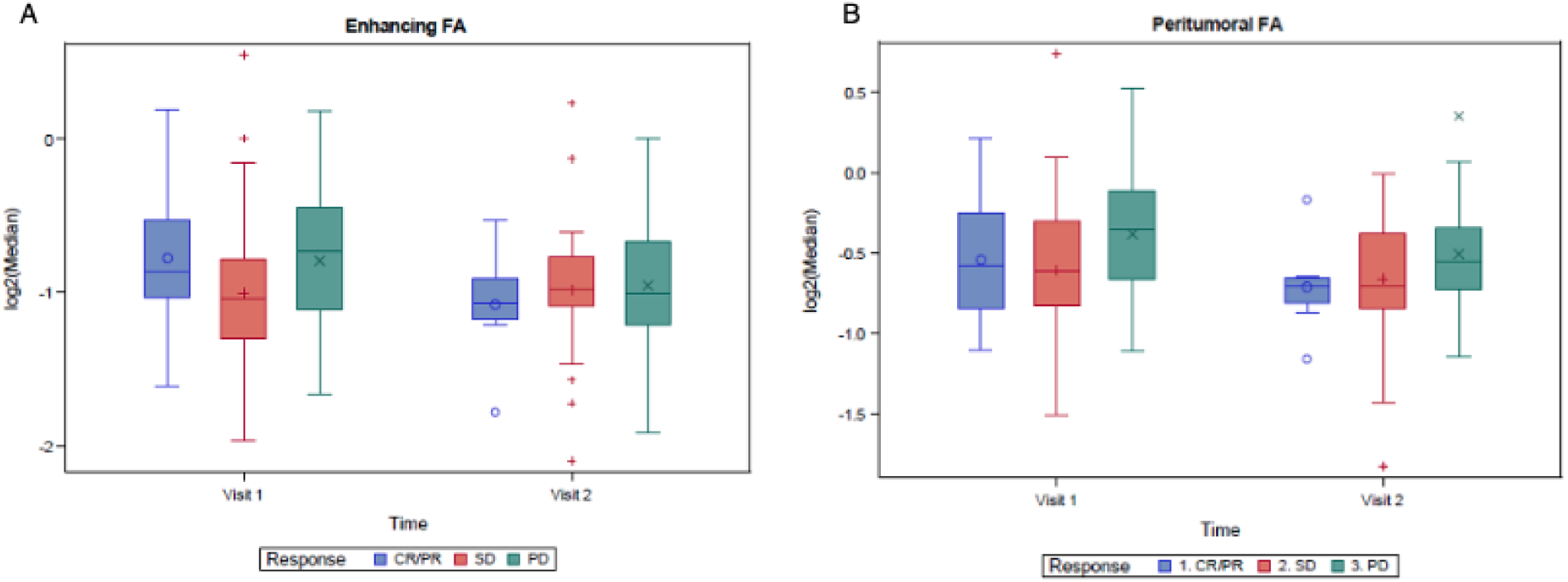
Fractional anisotropy within BM (A) and the peri-tumor region (B) during treatment with pembrolizumab, organized by response. There were no statistically significant differences in fractional anisotropy between response categories at either pre-treatment or post-treatment time points.

**Supplemental Figure 12:**
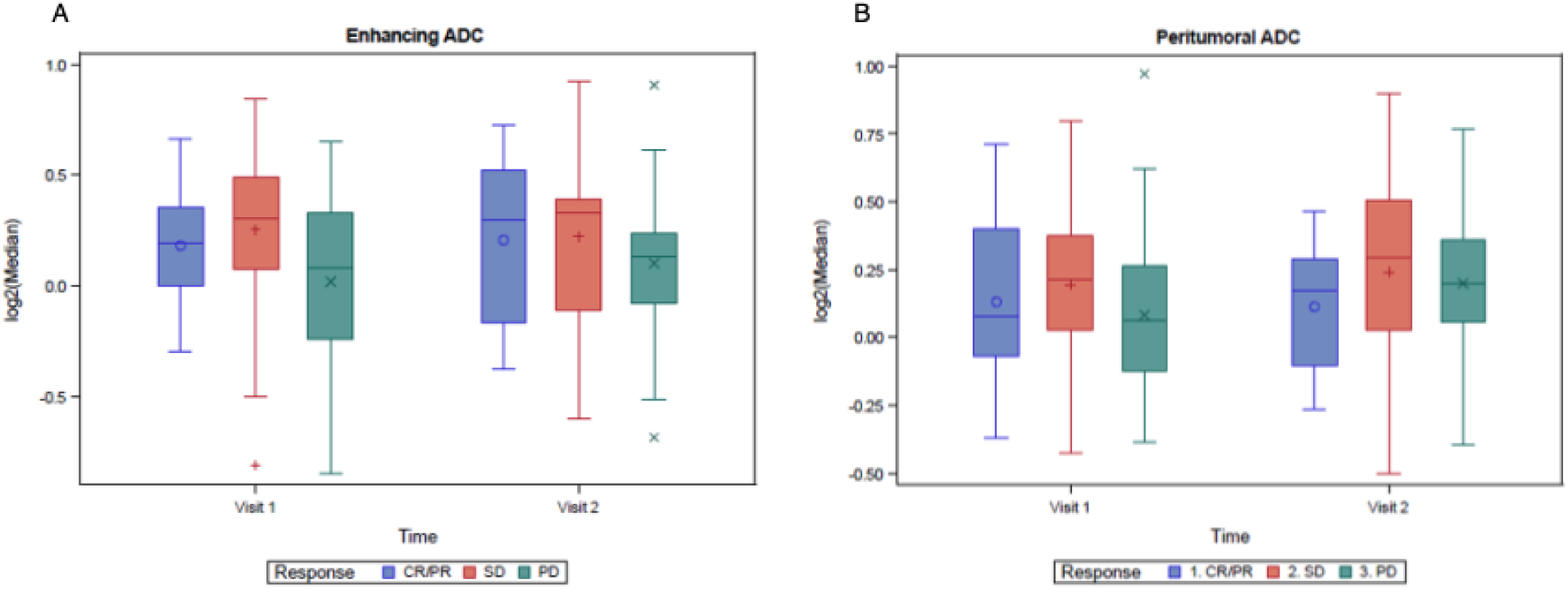
Apparent diffusion coefficient (ADC) within BM (A) and the peritumor region (B) during treatment with pembrolizumab, organized by response. There wa no statistically significant differences for ADC pre-treatment or post-treatment between response categories. PD BM displayed a statistically significant increase in median ADC for both the contrast-enhancing region (p = 0.04) and peri-tumor region (p = 0.006) between the pre-treatment and post-treatment visit. No significant differences were observed for BM with PR/CR (p = 0.85 – contrast enhancing and peri-tumor) or SD (p = 0.61 – contrast-enhancing; p = 0.93 – peri-tumor).

**Supplemental Table 1:** Aggregate metrics of vascular physiology. Pre-treatment and post-treatment columns reflect analysis of BM voxels before and after MR processing (e.g., voxels that met an adequate SNR) as defined in prior work^22,24^. The variable ‘Volume’ is the volume of the BM that was captured within the FOV of DSC MRI. MR processing is described in the Supplemental Methods and includes removal of image voxels of low quality.

|  | **Pre-treatment**<br>**(IQR)** | **Pre-treatment**<br>**after MR**<br>**processing (IQR)** | **Post-treatment**<br>**(IQR)** | **Post-treatment after**<br>**MR processing**<br>**(IQR)** |
| --- | --- | --- | --- | --- |
| **Volume (cm<sup>3</sup>)** | 2.19 (0.72 – 7.51) | 1.85 (0.60 – 5.28) | 2.82 (0.89 – 7.10) | 2.32 (0.80 – 6.53) |
| **Underperfused Tissue**<br>**(percentage)** | 12.43 (4.65 – 28.02) | 8.12 (2.56 – 18.93) | 8.12 (2.64 – 22.66) | 5.46 (0.86 – 15.57) |
| **Microvasculature**<br>**(percentage)** | 11.47 (3.73 – 26.00) | 9.73 (3.08 – 19.63) | 10.23 (3.25 – 21.29) | 6.94 (2.23 – 15.38) |
| **Arteries (percentage)** | 16.70 (10.60 – 25.35) | 20.12 (14.24 – 29.66) | 17.72 (12.49 – 26.32) | 22.95 (14.64 – 30.67) |
| **Veins (percentage)** | 20.73 (14.18 – 28.95) | 26.35 (18.20 – 34.50) | 21.95 (15.42 – 36.56) | 29.75 (19.87 – 40.67) |

**Supplemental Table 2:** Assessment of inter-patient heterogeneity of BM vascular physiology. For the 17 patients that possessed 3 or more BM at baseline, 11 patients had a breast primary, 1 patient had a lung primary, and 5 patients had primary tumors of other histologies. The median number of BM per patient was 3 (range: 3-5). For each vascular parameter, this analysis used the individual BM mean and ROI data to calculate the coefficient of variation (CV) for each BM at baseline. The p-value assessed if there were differences in average CV between histology. All vascular biomarkers for BM of the same patient were significantly more homogenous to each other, compared to those for BM of different patients.

| **Vascular biomarker** | **Patient p-value** |
| --- | --- |
| Vessel Size | <0.0001 |
| SE CBV | <0.0001 |
| GE CBV | 0.02 |
| K-Trans | 0.006 |

**Supplemental Table 3:**
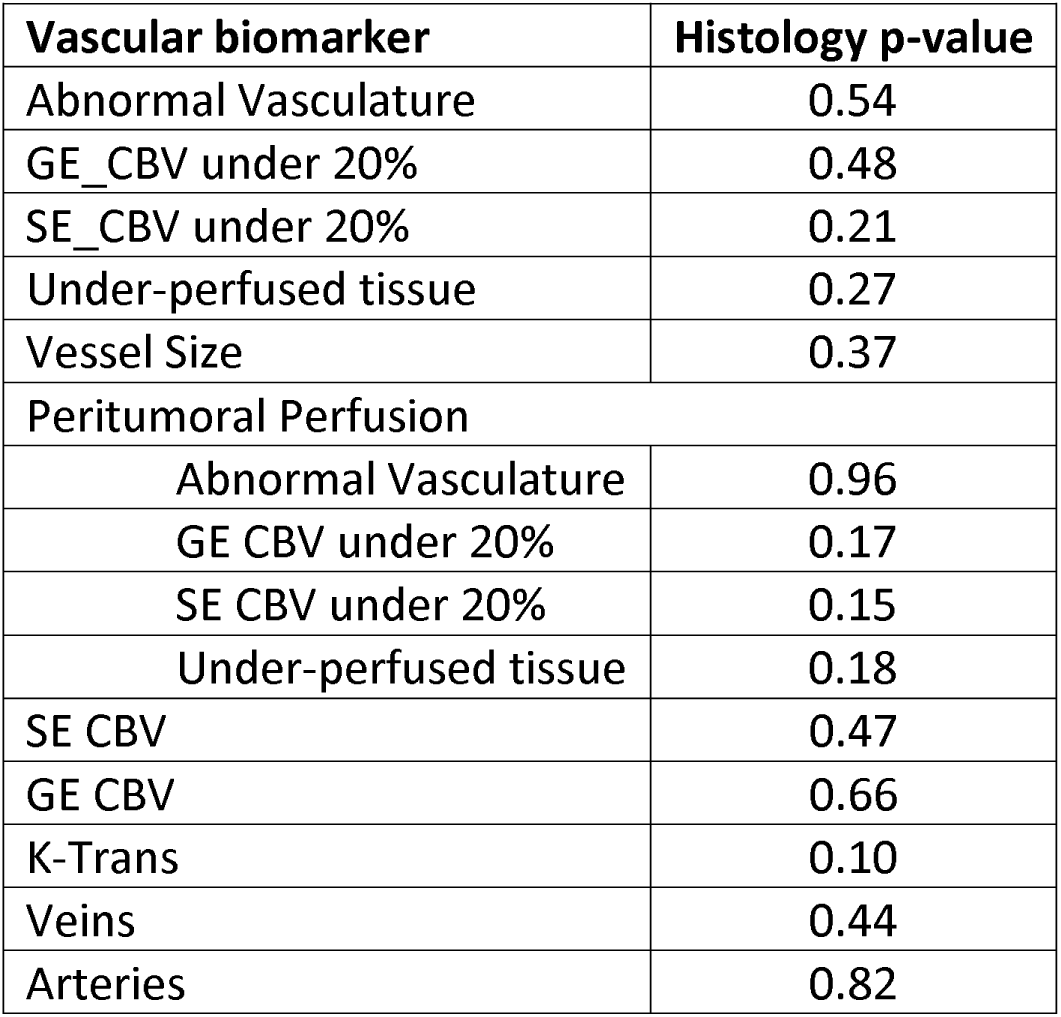
Assessment of inter-histology heterogeneity of BM vascular physiology. For this analysis, BM were classified according to histology: breast, lung, and other. The p-values reported below are for the effect of histology on the vascular biomarker of interest. There was no significant relationship noted between any vascular biomarker and histology.

## Supplemental Methods

### Research MRI acquisition protocol

T1-pre and T1-post contrast 2D axial images were obtained with TR 250 ms, TE 4.58 ms, FA 90°, slice thickness 5 mm, slice gap 1 mm, 25 slices, FOV 256 mm X 224 mm, and matrix size 256 X 256. T2SPACE 3D images were acquired with TR 3200 ms, TE 406 ms, echo train length 926ms with variable flip angles, slice thickness 1 mm, 176 slices, FOV 256 mm X 256 mm, and matrix size 256 X 256. MPRAGE 3D images were obtained with TR 2530 ms, 4 echoes with maximum TE 7.17ms, slice thickness 1 mm, 192 slices, FOV 270 mm X 256 mm, and matrix size 256 X 256.

T1 mapping was performed by repeating a T1-weighted 3D gradient-echo acquisition six times, each at a different flip angle (FA). The imaging parameters were TR 5.6 ms, TE 2.57 ms, slice thickness 3 mm, 36 slices, FOV 256 mm X 208 mm, matrix size 128 X 87, acceleration factor of 3 and 4 averages. The six FAs were 2°, 5°, 10°, 15°, 20°, and 30°. Following the T1 mapping, DCE-MRI was acquired by using a similar protocol as T1 mapping, except for 10° FA and 1 average, and 250 measurements (frames) collected in 412s, corresponding to a temporal resolution of 1.65 s. A bolus of 0.1 mmol/kg of gadoterate meglumine (Dotarem^®^, Guerbet USA) was injected 1 min after the DCE scan started, followed by 20 mL saline flush at 3.5 mL/s injection rate.

DSC-MRI was acquired by using a 2D echo-planer sequence with TR 1500 ms, TE 31 ms, FA 90°, slice thickness 5 mm, slice gap 1.5 mm, 11 slices, FOV 220 mm X 220 mm, and matrix size 128 X 128. A total of 99 frames were collected in 161 s, corresponding to a temporal resolution of 1.63 s. A bolus of 0.1 mmol/kg of gadoterate meglumine was injected 30 seconds after the DSC scan started, followed by 20 mL saline flush at 3.5 mL/s injection rate. The gadoterate meglumine injection for DCE-MRI served as preload for the DSC-MRI.

DCE-MRI and DSC-MRI data were motion corrected to align all volumes to the first volume of each dynamic series by using FSL.^2^ A deep learning-based model^3^ was used to automatically segment tumor region of interest (ROI) on the T1W post-contrast (corresponding to the CE volume). ROIs were visually inspected and manually corrected for accuracy. Diffusion and perfusion maps were generated within the CE volume. Maps of relative cerebral blood flow (rCBF) and volume (rCBV) to normal apparent grey and white matter tissue were obtained by using NordicICE (NordicNeuroLab AS, Bergen, Norway) with T1 leakage correction and population-based AIF.^4,5^ Permeability transfer constant (K^trans^) maps were generated from motion corrected DCE-MRI data by using custom software in MATLAB based on Tofts Model with a constant T1 of 1000 ms and population-based AIF.^6,7^

A 1-channel transmit combined with either an 8- or 32-channel receive radiofrequency coil array built to minimize 511 keV photon attenuation were used for the study.

### Additional image processing techniques

Co-registration of all MR sequences to the DSC space was performed using normalized mutual information in nordicICE. MRI perfusion analysis included motion correction, automatic detection of the arterial input function with deconvolution by standard single value decomposition, and contrast agent leakage-correction adapted for both T1- and T2-shortening effects^66^. We removed image voxels of low quality within the region-of-interest (ROI) by excluding voxels that were below the second percentile for signal-to-noise ratio (SNR) within the GE or SE maps across our cohort. First, using DSC GE or SE signal over time, the R2 curve was generated using:

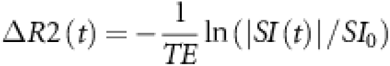

TE represented echo time (31 ms for GE; 104 ms for SE) and SI_0_ was the baseline DSC signal intensity at study state. Next, CNR was calculated using:

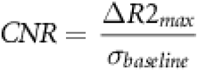

R2_max_ was the maximum R2 value over time, and σ_baseline_ was the standard deviation of the R2 curve during the baseline steady state after contrast administration. After removing voxels of low quality, the revised ROIs were overlaid on the MR sequence to derive the metric of interest.

## Acknowledgments

We thank the patients and their families for contributing to research efforts.

## Data Sharing Statement

Any requests for raw and analyzed data will be reviewed by the DF/HCC Institutional Review Board (IRB). Patient-related data not included in the paper were generated as part of a clinical trial and are subject to patient confidentiality. Any data and materials (e.g. tissue samples or imaging data) that can be shared will need approval from the DF/HCC IRB and a Material Transfer Agreement in place. All data shared will be deidentified.

## Code Availability

No code was used in this study

## Author Contributions

AEK, PKB, KEE, and ERG conceived the study. AEK and KWL performed data analysis, with input from KC, MG, KH, JBP, PS, CPB, SRA, BAB, WL, and EF-G, BRR, YY, and JK-C. AEK, PKB, EQL, NUL, BO, PYW, LN, JVC, JD, AE, RSH, IK, DL, EM, EW, MM, KO, HAS, DPC, ERG, and RJS supported the accompanying clinical trial, including recruitment of patients in the trial that underwent DSC-MRI. AG-H performed the statistical analysis. AEK, KWL, KEE and ERG wrote the manuscript. All the authors interpreted the data, reviewed the manuscript and approved the final version.

## Competing Interests Statement

PKB has consulted for Tesaro, Angiochem, Genentech-Roche, ElevateBio, Eli Lilly, SK Life Sciences, Pfizer, Voyager Therapeutics, Sintetica, and Dantari, received institutional research funding (to MGH) from Merck, Mirati, Eli Lilly, BMS and Pfizer and has received honoraria from Merck, Pfizer, and Genentech-Roche. BO has received clinical trial support from Incyte and Eisai. JD has served as a consultant for Amgen, Blue Earth Diagnostics and Unum Therapeutics and received research support (to MGH) from Novartis and Eli Lilly. RSH has consulted for AbbVie, Daichii Sankyo, EMD Serono, Lilly, Novartis, Regeneron, and Sanofi; and received institutional research funding (to MGH) from Abbvie, Agios, Corvus, Daichii Sankyo, Erasca, Exelixis, Lilly, Mirati, Novartis, and Turning Point. HAS has served on the scientific advisory board for Advanced Accelerator Applications, and received institutional research funding (to MGH) from AbbVie. RJS has consulted for Novartis, BMS, and Pfizer, and received institutional research funding (to MGH) from Merck.

## Notes

**Acknowledgments of Research Support** AEK is supported by an American Brain Tumor Association Basic Research Fellowship In Honor of Paul Fabbri, the William G. Kaelin, Jr., M.D., Physician-Scientist Award of the Damon Runyon Cancer Research Foundation, American Association for Cancer Research Breast Cancer Research Fellowship, and American Society of Clinical Oncology Young Investigator Award. EFG was supported by the European Union’s Horizon 2020 Programme: Marie SkłodowskalJCurie grant agreement No 844646-GLIOHAB and Spanish State Research Agency, Subprogram for Knowledge Generation (PROGRESS, No PID2021-127110OA-I00). PKB is supported by the NCI (1R01CA227156-01, 5R21CA220253-02 and 1R01CA244975-01), Damon Runyon Cancer Research Foundation, and Breast Cancer Research Foundation. YY is supported by the NIH (1R21AG067562-01 and 1R21GM137227-01). KEE is supported by the European Union’s Horizon 2020 Programme: ERC Grant Agreement No. 758657-ImPRESS, South-Eastern Norway Regional Health Authority (Grant Agreements No. 2016102, 2017073, 2013069), and the Norwegian Cancer Society and the Research Council of Norway FRIPRO Grant Agreements No. 261984, 303249). ERG is supported by R01CA211238-01 and R01CA244975.

